# STARgrid – an open end-to-end pipeline for Visium HD

**DOI:** 10.64898/2026.10.09.757882

**Authors:** Ling-Hong Hung, Ka Yee Yeung

## Abstract

Grid-based spatial transcriptomics quantifies expression by counting reads, assigning them to genes and, from their barcodes, to capture locations on a grid, which is registered to a microscope image. For Visium HD, only the vendor’s Space Ranger performs all these steps, and its licences prohibit redistributing it or software derived from it. We present STARgrid, an open-source, end-to-end, agent-ready pipeline from Visium HD FASTQ files and images to count matrices registered to the tissue. Its counts agree closely with Space Ranger’s (gene-total correlations at least 0.9989); it runs 2.0–3.4 times faster and counts 1.9–7.6% more molecules. Unlike Space Ranger, STARgrid always assigns each read to the closest-matching barcode, which gives 0.8–1.1% of matched reads a different location, mostly where adjacent-row barcodes overlap, and places another 2.9–4.6%. This choice and settling ties for each molecule from all its reads are the largest source of added molecules. Around nuclei, counts follow a reproducible pattern: more molecules in a ring just around each nucleus, usually with a dip under it. STARgrid registers the image by matching the nuclei to the whole pattern with the correlation ratio, even where the dip is weak or absent. On lung tissue imaged by Xenium on the same section, this places the nuclei where Xenium does or one capture location away. The added molecules behave as signal: they are inside cells almost as often as the others, and bring 15 of 20 abundant cancer genes closer to Xenium on an adjacent section (*P* = 0.0414).

## Introduction

Spatial transcriptomics measures where genes are expressed in a tissue section, so that expression can be assigned to cells and to larger structures such as tumour regions. Imaging methods hybridise fluorescent probes to a tissue section and read them under a microscope, recording position directly (Chen et al. 2015). Their gene panels can now comprise thousands of genes as in Xenium (Ozirmak Lermi et al. 2025; Janesick et al. 2023), up to the scale of the transcriptome (Eng et al. 2019; 10x Genomics 2026a). Capture methods take an alternative approach: they place the tissue section on a surface carrying a grid of positional barcodes, sequence the captured transcripts together with their barcodes, use the barcodes to place each transcript on the grid, and then align the grid with an image of the section, a step called registration. The first capture method, named Spatial Transcriptomics, used printed spots (Ståhl et al. 2016); later platforms reached finer resolution with beads (Rodriques et al. 2019), flow-cell clusters (Cho et al. 2021) and DNA nanoballs (Chen et al. 2022), and these platforms have been compared systematically (You et al. 2024).

As imaging panels grow, the difference between the two approaches has become less about the number of genes assayed and more about how position is obtained and how molecules are counted. In imaging methods, the fluorescent signal of the probes bound to a transcript identifies its gene and gives its position, in the same images that show the nuclei and boundaries of the cells. Capture methods make these measurements separately: a molecule’s position is read from its barcode, its identity from the rest of the read, and the structures of the tissue from a separate image of the section. Their resolution depends on the spacing of the capture grid and on how far molecules move before they are captured. At present capture methods are generally coarser than imaging, but in theory they are not limited by the optical resolution needed to separate two neighbouring fluorophores. In imaging that limit causes crowding: where transcripts are dense, their signals overlap and cannot be told apart, which caps how many can be detected (Eng et al. 2019). Furthermore, capture methods quantify expression by counting reads, which, as in RNA-seq, can give a much higher dynamic range than methods based on fluorescence measurements (Wang et al. 2009; Zhao et al. 2014).

In capture methods, the position of a read is found by matching the barcode it carries to a list of known barcodes, each of which maps to a position on the grid; this step is called barcode decoding. In Visium HD (de Oliveira et al. 2025) the barcoded surface is a continuous lawn of 2 µm squares. Each square is a capture location, identified by two coordinate barcodes, one for its column and one for its row, and counts are reported for each capture location and for 8 and 16 µm bins of neighbouring locations. Because of sequencing errors, the barcode in a read often differs from every listed barcode by a few substituted, missing or extra bases, so it has to be matched to a listed barcode that is similar but not identical. A read can also be equally similar to more than one capture location, and this tie must be resolved. Which of the similar barcodes a pipeline accepts, and how it resolves ties, decides where these reads are placed.

Each read must also be assigned to a gene, and the reads of each molecule counted as one. Visium HD has two methods of quantitation, both taken from 10x Genomics’ single-cell assays. RNA-seq quantitation captures transcripts by their poly(A) tails and sequences them, as in droplet single-cell RNA sequencing (Chromium 3*^′^*; Zheng et al. 2017), and its reads are assigned to genes by alignment to the genome. Flex quantitation hybridises pairs of probes to their target transcripts, ligates each pair and sequences the ligated probes, as in Chromium Flex, and its reads are assigned to genes by matching the probe sequences. Each molecule is tagged on capture with a unique molecular identifier (UMI), a short random sequence, so reads of the same gene with the same barcode and the same UMI, or a UMI that differs by one base, are treated as copies of one molecule. In single-cell data the barcode identifies a cell; in Visium HD it identifies a capture location.

The last step is to register the microscope image of the section to the capture grid, so that the counts can be related to nuclei, cells and tissue structures. Assigning transcripts to cells has been called a central unresolved problem of spatial omics (Ishaque et al. 2026), and it depends on registration: an offset of a few micrometres is enough to move capture locations into or out of nuclei. In Visium HD, the section is imaged under the microscope on its own slide and then aligned with the capture slide in the CytAssist instrument, which records a second image that shows the capture slide’s fiducial marks. Space Ranger registers the microscope image to the grid through this image and its fiducial marks and, starting in version 4.0, refines the registration using transcript counts (10x Genomics 2026d). Besides automated checks for a failed registration, its documentation recommends inspecting overlays of the image and the counts. Open tools that start from Space Ranger’s output, such as Bin2cell (Polański et al. 2024), use its registration, and spacemake aligns an H&E image to an image built from the counts (Rajewsky lab 2026). However, we found no documented open method that uses the counts to register the image in each region of a slide and has been tested against an independent reference.

Space Ranger (10x Genomics 2026b) is currently the only pipeline that performs all four steps for Visium HD. None of the open tools we reviewed (Kijima et al. 2023; Sztanka-Toth et al. 2022; Xu et al. 2025; Vaidya et al. 2026) implements barcode decoding, gene assignment, count assignment to capture locations and bins, and registration (Supplemental Table S1). Space Ranger’s end-user licence prohibits modifying it, creating derivative software and reverse engineering it (10x Genomics 2026f). Its published source may be modified only for internal use and may not be redistributed, and neither may software derived from it (10x Genomics 2026e). Testing, comparing or improving these steps in the open therefore requires an open and independent implementation.

We built one on STAR Suite (Hung et al. 2026), our extension of the aligner STAR (Dobin et al. 2013) and its single-cell counter STARsolo (Kaminow et al. 2021). STAR Suite is a single executable program: one command takes an assay from sequencing reads to count matrices, with no chain of separate tools to install and connect. Its modules process bulk and single-cell RNA-seq (STAR-core), feature barcodes, as in CRISPR screens (STAR-perturb), probe-based Flex assays (STAR-Flex) and metabolic labelling of RNA (STAR-SLAM). Each assay has a recipe, which lists its settings and their permitted values. A person fills it in as a form in the browser-based Launchpad, and an AI agent through a Model Context Protocol server; both give the same command. These modules already assigned reads to genes and counted the molecules of each cell. None of them handled spatial barcodes or images.

Here we present STARgrid, an open-source, end-to-end and agent-ready pipeline for Visium HD, built in the same framework and released under the MIT licence. It goes from sequencing reads and images to count matrices registered to the tissue. We wrote it as a clean-room implementation, from the published documentation, the reference files distributed with Space Ranger and Space Ranger’s output, without reading or decompiling any 10x Genomics code, so that no part of it derives from Space Ranger and it can be redistributed (Methods). Only gene assignment is taken from STAR Suite; the other three steps are new. The first is the decoding of spatial barcodes: STARgrid always assigns a read to the capture locations whose barcodes match it most closely. The second is count assignment, with its handling of ambiguity: when several locations match equally well, it keeps them all and chooses among them only after the reads of each molecule have been grouped. Both are built into the STAR Suite program, so for either method of quantitation a single run of it decodes the spatial barcodes, assigns reads to genes and counts to capture locations, and writes matrices at 2, 8 and 16 µm. The third is registration, a separate step that follows this run and registers the image with the counts themselves. Around each nucleus, more molecules are counted in a ring just outside it and, usually, fewer under it. STARgrid moves the nuclei in the image until they best fit this pattern, for the whole slide and region by region. It measures the fit with the correlation ratio, a similarity measure taken from the registration of different kinds of medical images (Roche et al. 1998). Because the measure uses the whole pattern, it works even where nuclei do not hold fewer molecules than the tissue around them. The run and the registration each have a recipe, so a person or an agent can set them up in the same way.

We compare the output and running time of STARgrid with those of Space Ranger on public human slides that cover both methods of quantitation, examine individual reads to find where the two pipelines place them differently, and measure where the molecules that STARgrid adds come from. We also compare the registration with Space Ranger’s and test it in lung tissue against nuclei imaged by Xenium on the same section. Finally, we test whether the added molecules are signal: whether they lie inside cells in the slide’s own H&E image, and whether, for a panel of cancer genes, they bring the counts closer to Xenium RNA imaging of an adjacent section of an ovarian cancer.

## Results

### STARgrid agrees with Space Ranger, counts more molecules and runs faster

STARgrid computes the location and identity of each molecule in four steps (Fig. 1A,B): barcode decoding, gene assignment and count assignment in a single run, and then the registration of the image to the capture grid, which uses the images and the counts (Methods). Decoding, count assignment and registration are new; gene assignment uses the methods of STAR Suite (Hung et al. 2026). We compared the output with Space Ranger’s on four human slides (Table 1). Two are 10x Genomics demonstration slides of colorectal cancer with Flex quantitation: the colorectal slide (H1-GMHFWPH/D1) and slide H1-VM2JXXK/A1. The SPATCH slide is a Flex slide of ovarian cancer from SPATCH, a spatial transcriptomics resource and benchmarking framework (Ren et al. 2025), and the ovarian dataset a 10x Genomics demonstration slide of ovarian cancer with RNA-seq quantitation.

**Figure 1.**
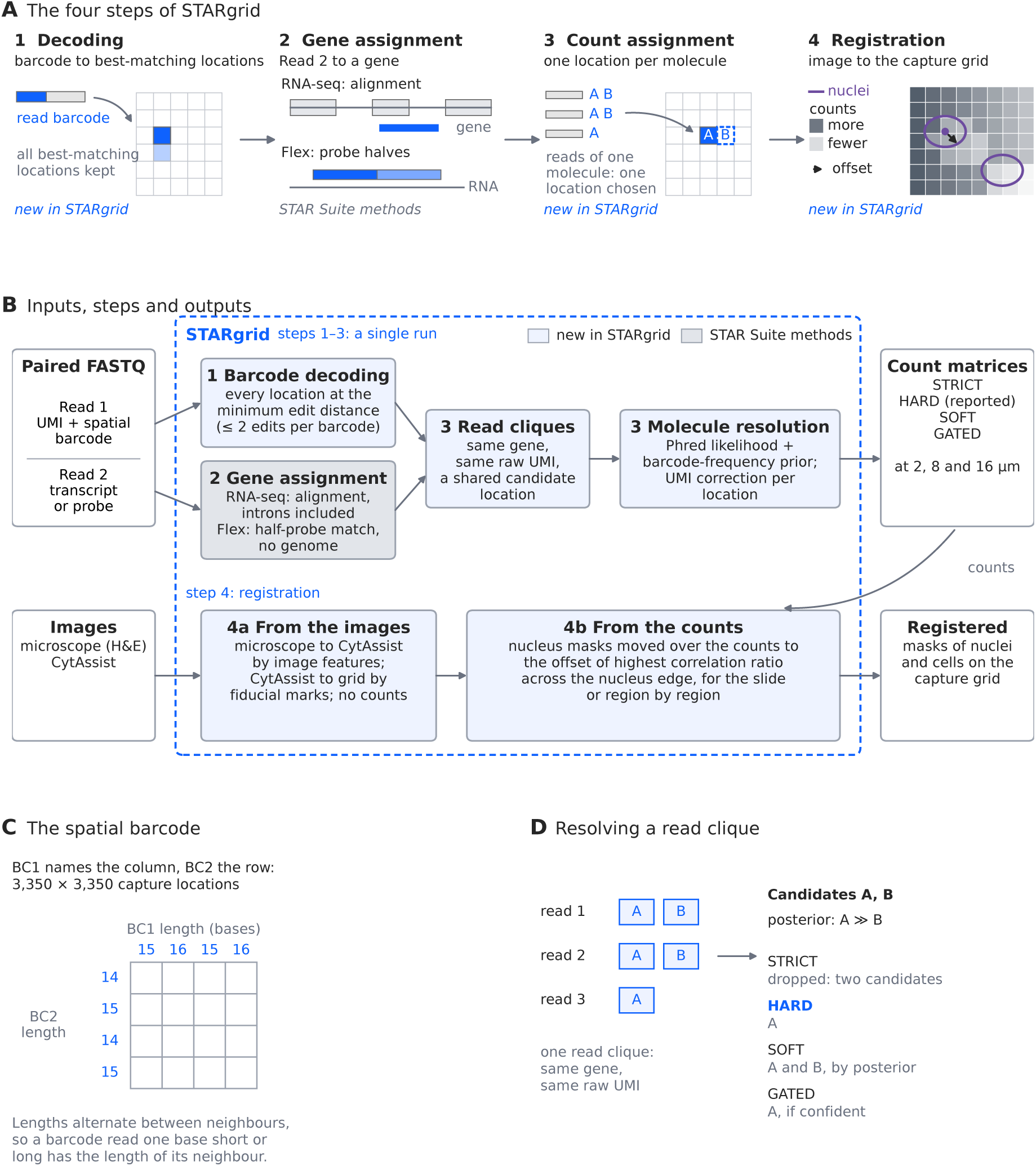
The four steps of STARgrid. **A**, STARgrid matches the spatial barcode of each read to the grid barcodes, keeping the best match and any ties (1); assigns Read 2 to a gene, by alignment to the genome for RNA-seq quantitation or by matching the halves of each probe for Flex quantitation (2); groups the reads of each molecule and chooses one location for the molecule from all of them (3); and registers the microscope image to the capture grid (4; Fig. 4D,E). Steps 1–3 take place in a single run of the STAR Suite program (Methods); step 4 uses the images and the counts. Steps 1, 3 and 4 are new in STARgrid (blue boxes in **B**); step 2 uses the methods of STAR Suite (grey box). **B**, Inputs, steps and outputs. Reads sharing a gene, a raw UMI and a candidate location form a read clique, which is resolved to one or more capture locations, or left out, depending on the resolution policy; count matrices are written at 2, 8 and 16 µm under each resolution policy. The registration has two stages: the first (4a) uses the microscope and CytAssist images and no counts; the second (4b) moves the nuclei in the image to the offset at which the correlation ratio of the counts is highest, for the whole slide or region by region, and so places the nucleus and cell masks on the capture grid. **C**, BC1 names the column and BC2 the row of the 3,350 *×* 3,350 grid. BC1 barcodes alternate between 15 and 16 bases across columns and BC2 barcodes between 14 and 15 across rows, so a barcode read one base short or long has the length of its neighbour. **D**, A location can be a candidate only within 2 edits in each barcode and the documented total of four. If one candidate has the fewest edits in total, summed over BC1 and BC2, only it is kept; if several tie, all of the tied candidates are kept. Candidates with more edits are dropped. Here three reads of one clique share candidates A and B; STARgrid assigns the clique to its most probable location (_HARD_), and the other policies shown can be applied to the same run. The posterior combines the reads’ log-likelihoods, from the quality scores of their barcode bases, with a barcode-frequency prior estimated in each run from the reads that match a single location with no edits.

**Table 1.** Slides and results at a glance. Four slides were compared with Space Ranger; the first three were also used to time the two pipelines and to test the counts that STARgrid adds against the H&E image of the same section. Differences are 100 *×* (STARgrid *−* SR)*/*SR. Wall times are medians of three runs of each pipeline on the same computer, each with a cold file cache, in minutes:seconds; both include the registration of the image, which in STARgrid, unlike Space Ranger, includes segmenting the nuclei on a GPU (Methods). On slide H1-VM2JXXK/A1, Space Ranger 4.0.1 (reference GRCh38-2024-A) was run with BAM output, which records the location of each read, for the read-by-read comparison (Table 2). The lung tissue that tested the registration is listed with the other datasets in Supplemental Table S3. A dash marks a measure not made on that slide.

|  | Colorectal (Flex) | SPATCH ovarian (Flex) | Ovarian (RNA-seq) | Colorectal, H1-VM2JXXK/A1 (Flex) |
|---|---|---|---|---|
| Slide | H1-GMHFWPH/D1 | H1-WP2FTZD/D1 | H1-YQJBZ7X/D1 | H1-VM2JXXK/A1 |
| Tissue | colorectal cancer, FFPE | ovarian cancer, FFPE | ovarian cancer, fresh frozen | colorectal cancer, FFPE |
| Source | 10x Genomics | SPATCH | 10x Genomics | 10x Genomics; GEO |
| Published with | FASTQ, images, SR 4.1.0 output | reads, images, SR filtered matrix | FASTQ, images, SR 4.0.1 output | FASTQ, SR output |
| Probe set or genome reference | 2020-A | 2020-A | 2024-A | 2024-A |
| Read pairs | 212,554,625 | 676,879,511 | 474,131,092 | 721,219,716 |
| Space Ranger comparator | published 4.1.0, raw | published, filtered | our 4.1.0 run, introns, raw | our 4.0.1 run, BAM, raw |
| Further references | H&E | H&E | H&E; Xenium | Space Ranger’s read locations |
| *Agreement with Space Ranger* |  |  |  |  |
| Gene-total Pearson *r* | 0.999999 | 0.999997 | 0.9989 | 0.9999997 |
| Bin-total Pearson *r* | 0.99980 (16 µm) | 0.99877 (8 µm) | 0.9994 (8 µm) | 0.99981 (16 µm) |
| *Molecules* |  |  |  |  |
| Space Ranger | 72,861,605 | 264,118,518 | 187,552,358 | 272,612,275 |
| STARgrid | 76,226,984 | 270,244,052 | 201,888,275 | 279,777,220 |
| Difference | +4.62% | +2.32% | +7.64% | +2.63% |
| Difference, bins common to both outputs | +4.62% | +1.92% | +7.64% | +2.63% |
| *Wall time, FASTQ files and images to registered counts* |  |  |  |  |
| STARgrid / Space Ranger | 9:41 / 26:00 | 25:21 / 86:27 | 30:31 / 59:42 | — |
| Times faster | 2.7 | 3.4 | 2.0 | — |
| *Where the added counts fall (H&E of the same section)* |  |  |  |  |
| Cell ROC AUC of all counts, STARgrid minus Space Ranger | 0.000079 | 0.000055 | — | — |
| In-cell fraction, added minus shared (points) | −1.15 | −0.93 | −0.86 | — |

The matrices agree closely with Space Ranger’s (Fig. 2; Table 1; full comparison in Supplemental Table S2). However, in the bins common to both outputs, STARgrid counts more molecules than Space Ranger on every slide: 4.62% more on the colorectal slide, 2.63% on slide H1-VM2JXXK/A1, 1.92% on the SPATCH slide and 7.64% on the ovarian dataset.

**Figure 2.**
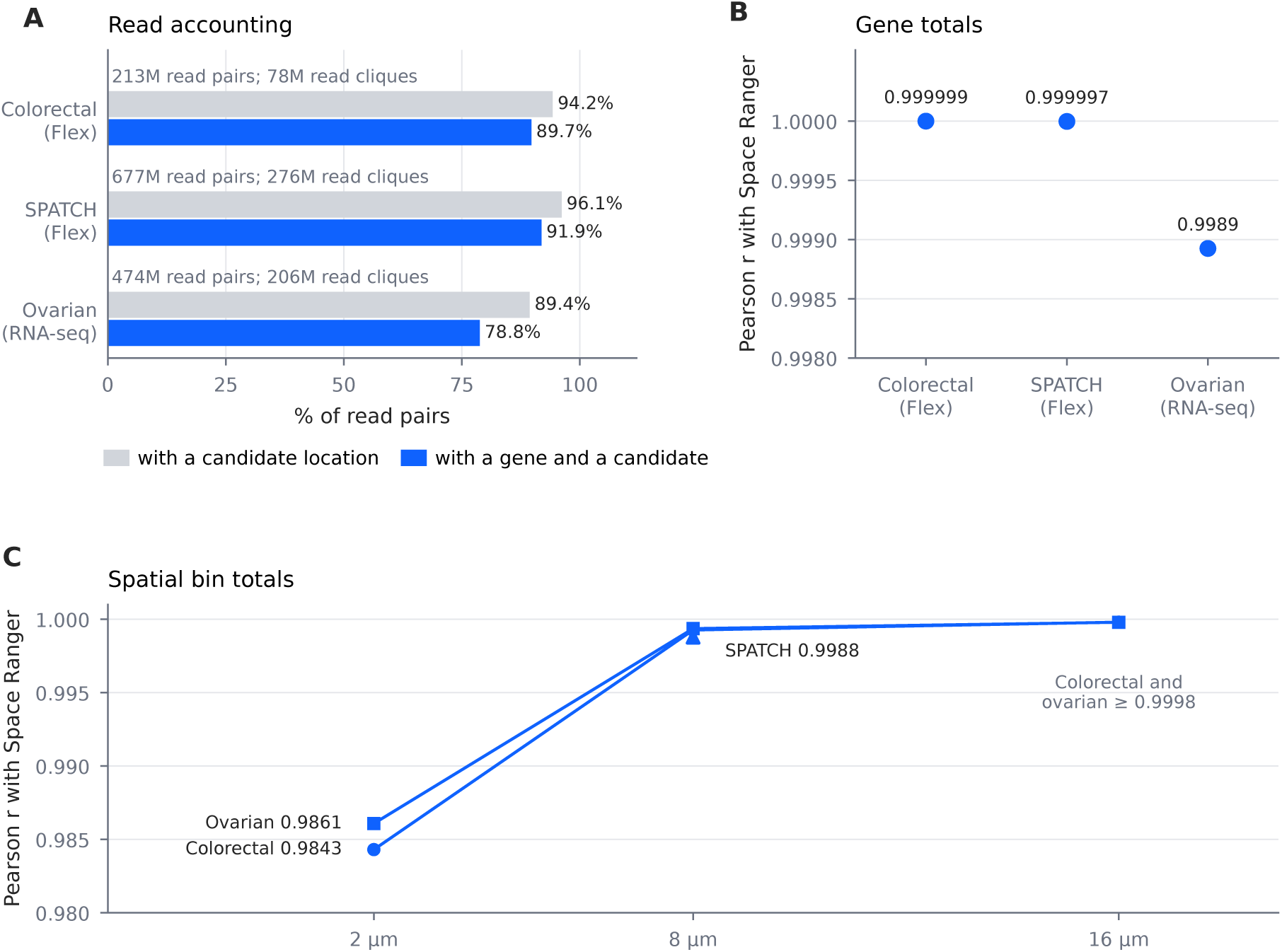
The output largely agrees with Space Ranger’s on three slides, with both Flex and RNA-seq quantitation. **A**, Read accounting: the share of read pairs with a candidate location, and with both a gene and a candidate location, with the numbers of read pairs and read cliques. **B**, Pearson correlation of gene totals with Space Ranger’s. **C**, Pearson correlation of spatial bin totals at 2, 8 and 16 µm over the whole capture grid (colorectal, ovarian), and at 8 µm over the bins of the published filtered matrix (SPATCH). STARgrid’s counts are those under _HARD_, and the ovarian comparator is Space Ranger 4.1.0 run with introns included.

With 32 processor threads each, STARgrid converts the FASTQ files into counts registered to the image 2.0 to 3.4 times faster than Space Ranger (Table 1). This includes work that Space Ranger’s runs did not do: because STARgrid’s registration fits the nuclei to the pattern of counts around them, it also segmented the nuclei, on a GPU, while the reads were processed. Methods lists this and the other differences between the two configurations.

### Decoding and count assignment: STARgrid places more reads, always at their closest-matching barcodes, and this is the largest source of the added molecules

Each read pair has two reads: Read 1 carries the UMI followed by the two coordinate barcodes of the capture location, and Read 2 the sequence of the transcript or probe (Fig. 1B). We call the coordinate barcode of the column BC1 and that of the row BC2; the same two lists of barcodes are used on all the slides analysed here. There are 3,350 of each, and so 11,222,500 capture locations (Fig. 1C). The barcodes are not all the same length: column barcodes are 15 or 16 bases and row barcodes 14 or 15. Neighbouring columns have column barcodes of different lengths, and so do neighbouring rows. Where the column barcode ends and the row barcode begins in a read therefore depends on the location the read came from, and is not known until the read has been matched. As a result, a read with a single missing or extra base can match the coordinate barcodes of neighbouring locations equally well. Ambiguity can also arise from the two lengths themselves: for 182 pairs of adjacent rows, the shorter row barcode is the longer one without its first base, so a read from one of these rows can also match the other (Methods).

To see where the two pipelines place reads differently, we compared them read by read on slide H1-VM2JXXK/A1 and the ovarian dataset, using the BAM file written by Space Ranger, which records the location of each read (Fig. 3; Table 2; Methods). For most reads, Space Ranger’s location is among the best matches that STARgrid finds. The two pipelines place reads differently for three reasons: Space Ranger’s choice between overlapping rows, the reads that Space Ranger leaves without a location, and reads that match more than one location equally well.

**Figure 3.**
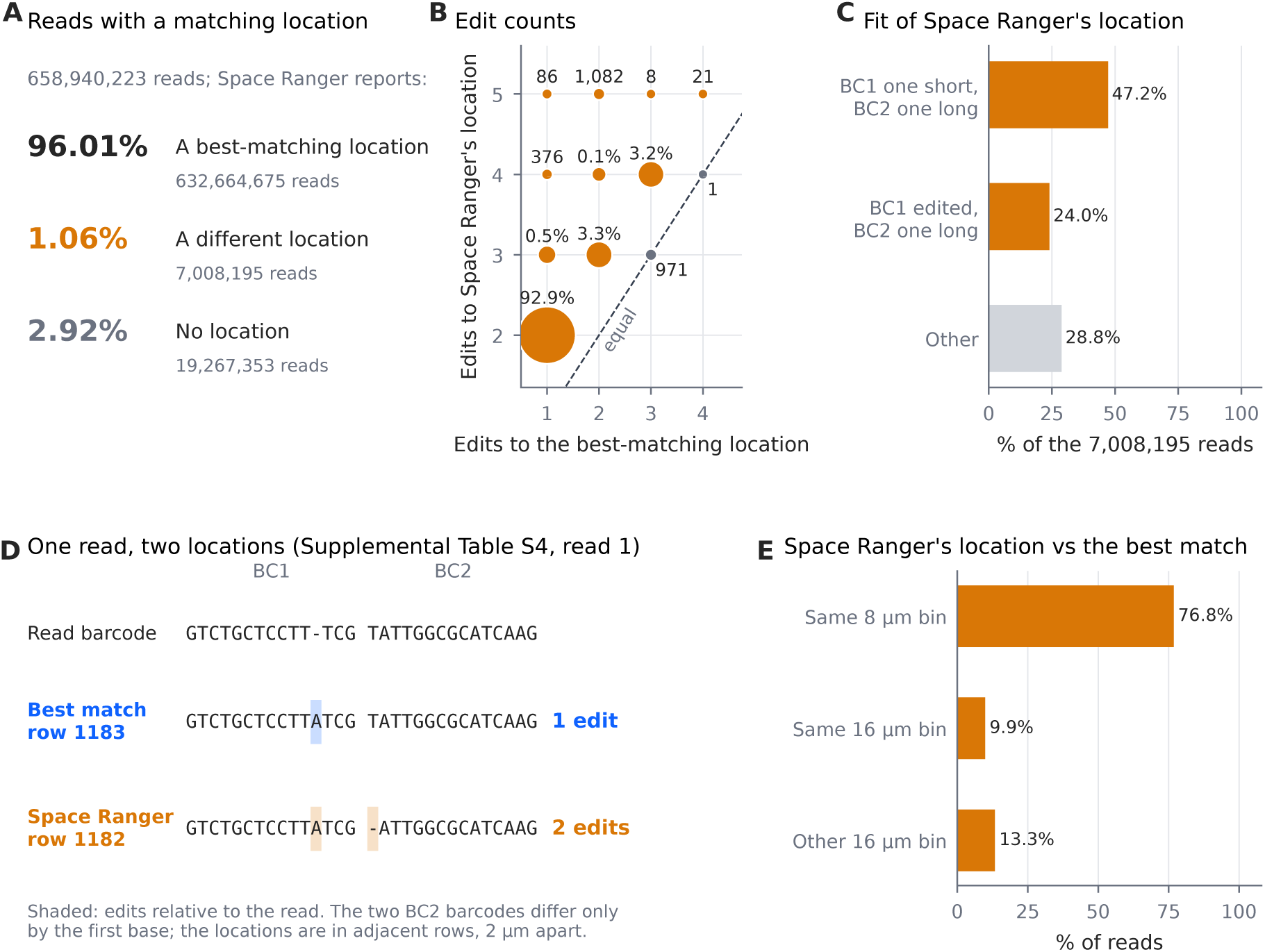
Space Ranger and STARgrid disagree on handling reads with overlapping barcodes. Colorectal slide H1-VM2JXXK/A1: the reads in the BAM file written by Space Ranger 4.0.1 that it assigned to a single gene and gave a UMI (Supplemental Methods). **A**, Of the 658,940,223 reads with a location within 2 edits in each barcode, the share for which Space Ranger’s location is a best-matching location, a different location, or absent. **B**, For the 7,008,195 reads with a different location: the edits between the read’s barcode and the best-matching location, against the edits to Space Ranger’s location. Edits are counted over every start of the barcode in Read 1 and every division of it into its two parts, with no limit on each barcode. Circle area is proportional to reads. No read lies below the diagonal. **C**, How the read’s barcode fits Space Ranger’s location: which barcode is read one base short or long, or edited. **D**, A worked read (Supplemental Table S4, read 1). The best-matching location needs one deletion in BC1; Space Ranger’s location, in the adjacent row, needs the same deletion and an inserted base in BC2. **E**, Where Space Ranger’s location lies relative to the best-matching location. The panels describe Space Ranger’s recorded output, not how it is produced.

**Table 2.** Where STARgrid and Space Ranger differ, read by read. Reads in the BAM file that Space Ranger wrote, compared with the capture locations that STARgrid finds for them (Fig. 3; Methods). Percentages in the second row are of the reads in the first; those under “Reads with a matching location” are of those reads, and indented rows are percentages of the row above. The molecules placed differently are those matched between the two pipelines by their reads. The difference in molecule totals is divided into three components, as percentages of Space Ranger’s total (Methods; counts in Supplemental Table S6); the components add up to the total, up to rounding.

|  | Colorectal,<br>H1-VM2JXXK/A1<br>(Flex) | Ovarian<br>(RNA-seq) |
|---|---|---|
| *Reads* |  |  |
| Assigned by Space Ranger to a single gene, with a UMI | 707,409,359 | 392,509,264 |
| With a location matching within STARgrid’s limits (%) | 93.15 | 91.62 |
| *Reads with a matching location (%)* |  |  |
| Matching more than one location equally well | 22.7 | 18.7 |
| Space Ranger’s location among the best matches | 96.0 | 94.7 |
| Space Ranger’s location not among the best matches | 1.06 | 0.77 |
| in one of two adjacent rows whose barcodes overlap | 93.38 | 92.48 |
| Left without a location by Space Ranger | 2.92 | 4.56 |
| with a single best match | 53.0 | 56.0 |
| Placed by Space Ranger with no matching location (reads) | 92 | 6 |
| *Molecules placed differently, explained by overlapping rows (%)* |  |  |
| In different 2 µm locations | 20.55 | 9.70 |
| In different 16 µm bins | 9.03 | 6.31 |
| *Difference in molecule totals (% of Space Ranger’s total)* |  |  |
| Placement: STARgrid’s locations, with Space Ranger’s reads, genes and UMIs | +2.62 | +4.65 |
| Gene assignment: STARgrid’s genes, selection of reads and handling of UMIs with reads for two genes | −0.18 | +1.95 |
| Deduplication: STARgrid’s collapsing of reads into molecules by UMI, at Space Ranger’s locations and with its genes | +0.18 | +1.04 |
| Total | +2.63 | +7.64 |

First, most of the reads whose Space Ranger location was not one of the best matches could be assigned to either of two adjacent rows whose barcodes overlap. In those reads, Space Ranger’s location had the shorter row barcode, whereas STARgrid, as for every read, keeps the location whose barcodes match with the fewest edits (two such reads are shown in Supplemental Table S4). Because the two rows are adjacent, this policy difference moves a molecule by one row, and most of the molecules it moves stay in the same 16 µm bin.

Second, Space Ranger leaves some reads without a location although their barcodes match a location within STARgrid’s limits, and STARgrid places them. Some of these reads have a single best match; the others match more than one location equally well and are placed with the other reads of their molecule (below). These are the reads with the most divergent barcodes: almost all of them needed two or more edits, and the share that Space Ranger left out rose with the number of edits (Supplemental Table S5). The opposite case is rare: almost every read that Space Ranger places, STARgrid places too, so the molecules that Space Ranger locates are nearly a subset of those that STARgrid locates.

Third, many reads match more than one capture location equally well. Space Ranger determines the location of each read on its own (10x Genomics 2026c), whereas STARgrid keeps every best-matching location until the reads of a molecule have been grouped, and then assigns the molecule to its most probable location (Fig. 1D; Methods).

To find out how much of the difference in molecule totals comes from where the two pipelines place reads, we divided it into three components: placement, gene assignment and deduplication (Table 2; Methods; Supplemental Table S6). Placement, which combines the decoding differences above with the choice of location for each molecule, was the largest component on both slides: the molecules that STARgrid adds come mainly from where it places reads, not from how it assigns genes or collapses reads into molecules.

### Gene assignment: gene totals agree closely with Space Ranger’s

STARgrid assigns each read to a gene in the same run that decodes its barcode, with methods described previously (Hung et al. 2026): by alignment to the genome for RNA-seq quantitation and by matching the two halves of each probe for Flex quantitation (Methods). Gene totals agree closely with Space Ranger’s on every slide (Pearson *r* of at least 0.9989; Table 1).

### Registration: the profile of counts around nuclei registers the image to the grid

To relate the counts to the tissue, the image must be registered to the capture grid (Fig. 1A, step 4). Registration from the images and the slide’s fiducial marks alone was inconsistent: it was in register on the colorectal slide but 8.49 µm off on the ovarian dataset and 8.2–15.2 µm off across the SPATCH slide (Fig. 4B,C; Fig. 5A,B). We noticed a reproducible pattern in the counts around nuclei: more molecules in a ring just around each nucleus and, usually, fewer under the nucleus itself. Moving nucleus masks over the counts therefore traces a dip where the masks lie on the nuclei, and the pattern holds nucleus by nucleus (Fig. 4A–C; Supplemental Figs. S1, S2; Supplemental Table S7). The dip, however, was shallow or missing in some tissues, such as kidney and mouse brain (public slides, below), where the nuclei held more molecules than the tissue just around them. In every tissue, the counts changed with the distance from the nucleus edge (Fig. 4D).

**Figure 4.**
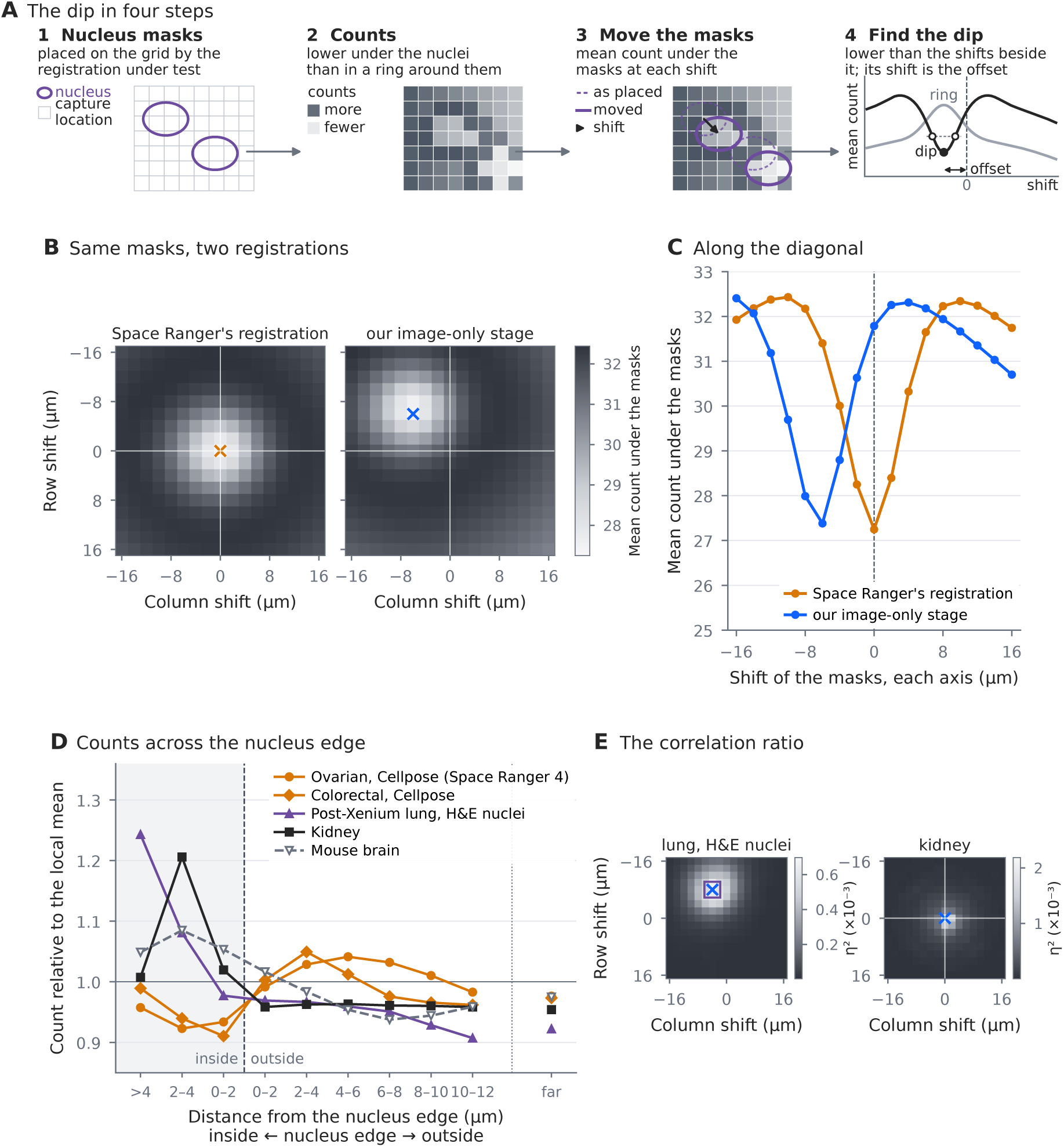
How the pattern of counts around nuclei registers the image. **A**, The dip, drawn schematically. Nucleus masks segmented from the microscope image are placed on the capture grid by a registration (1). On average, the count on the capture locations under the nuclei is lower than on a ring of locations just around them: a local hollow, which does not require the nuclei to hold fewer counts than the rest of the tissue (2; the hollow on the left holds more counts than the tissue on the right). The masks are moved over the grid location by location, and the mean count under them is recorded at each shift (3). The mean is lower where the masks lie on the nuclei than at the shifts one capture location around that position, a dip, and the shift of the deepest dip from zero estimates the offset (4). A mask of the ring, moved in the same way, has its highest mean at the same shift, a peak (4, grey). **B**, Mean count under our Cellpose nucleus masks of the ovarian dataset at each shift of up to 16 µm on each axis, with the masks placed by the registration in Space Ranger’s output deposited with the dataset (left) and by the image-only stage of our registration (right), on one grey scale (lighter, lower mean); STARgrid counts under _HARD_. Cross, the deepest dip; white lines, zero shift. Rows run down the slide. The window is chosen for display; the search used in each analysis is given in the Supplemental Methods. **C**, The same means along the diagonal of B, where the row and column shifts are equal, against the shift on each axis; dashed line, zero shift. **D**, The profile of counts across the nucleus edge on five whole slides, at the shift that the correlation ratio finds. Each point is the mean count in a band of distance from the edge of the nearest nucleus, inside the nuclei (shaded) or outside them, or in the rest of the tissue (far); each count is first divided by the mean count of the capture locations under tissue within 4 µm of it. Nuclei: our Cellpose-SAM masks placed by the Space Ranger version-4 registrations deposited with the ovarian dataset and the colorectal slide; the nuclei segmented from the H&E image of the post-Xenium lung section H1-9QZXXNP/D1, placed by the dataset’s registration; and Space Ranger’s nucleus maps of the public kidney and mouse brain slides. On the tumour slides the counts are lowest inside the nuclei near their edge and highest a few micrometres outside them, the dip and the ring; on the kidney and mouse brain slides they are highest inside the nuclei. **E**, The correlation ratio *η*^2^ at each shift of up to 16 µm on each axis, for the lung section (left) and the kidney slide (right) of D, each on its own grey scale (lighter, higher). Cross, the highest maximum; square, the shift that brings the H&E nuclei onto the nuclei imaged by Xenium on the same section; white lines, zero shift, which on the kidney slide is Space Ranger’s registration. Rows run down the slide.

**Figure 5.**
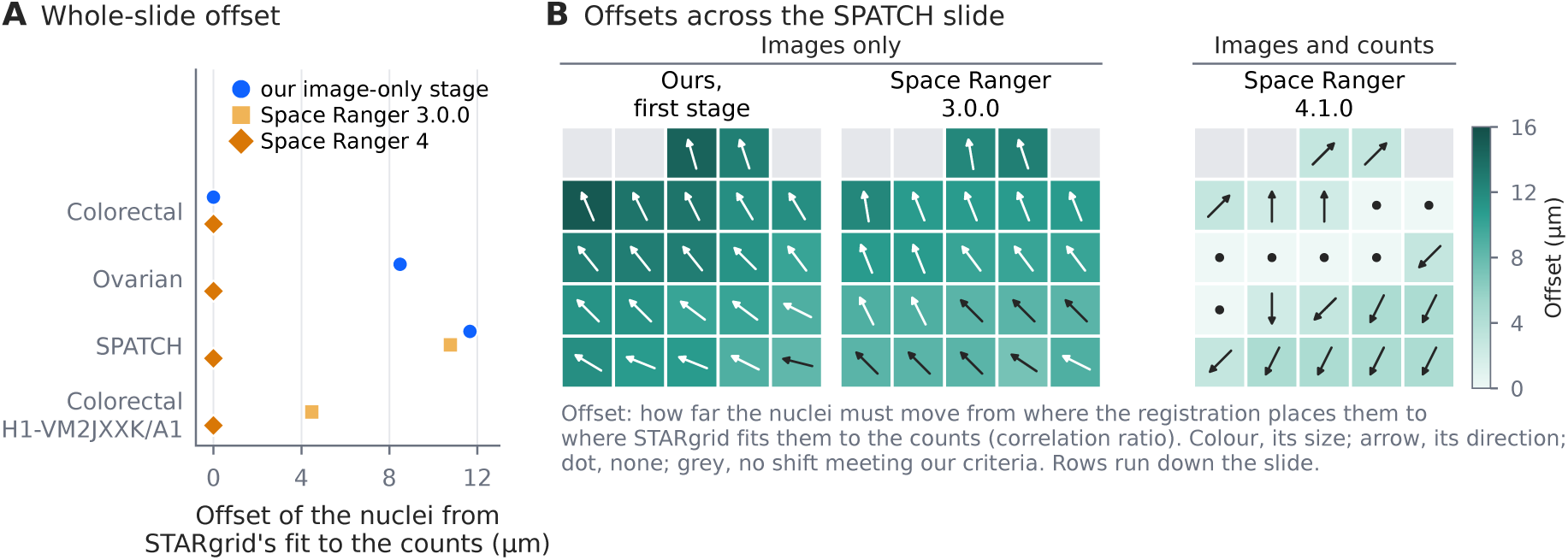
What registration with the counts found. **A**, Whole-slide offset under each registration: how far the nuclei must be moved from where the registration places them to where STARgrid fits them to the profile of counts around nuclei with the correlation ratio, to the nearest capture location. Zero means that the registration agrees with STARgrid’s fit. The nuclei are our nucleus masks, segmented with Cellpose-SAM; on the ovarian dataset and the two colorectal slides, masks segmented with StarDist, the control, give the same offsets (Supplemental Table S8). Ours is the first stage of our registration, which uses the images only. Space Ranger 4 is the registration in the output deposited with the dataset (ovarian, version 4.0.1; colorectal, version 4.1.0) or in our version 4.1.0 run on a subsample of 100,000 reads with the same image (SPATCH, H1-VM2JXXK/A1). Space Ranger 3.0.0 is our run on the same subsample with the same image (SPATCH) or the output deposited in GEO (H1-VM2JXXK/A1). **B**, The same offset in each region of a 5 *×* 5 division of the SPATCH slide. The registrations made from the images and fiducial marks alone, the first stage of ours and Space Ranger 3.0.0’s, leave large offsets across the slide. Space Ranger 4.1.0’s, which also uses the counts, by a different method from ours, comes close to STARgrid’s fit. The masks and the counts are the same in every map; only the registration differs. Colour, size of the offset, on one scale for every map; arrow, its direction; dot, no offset to the nearest location; grey, no shift meeting our criteria (Supplemental Methods). Rows run down the slide. Values for A and B are in Supplemental Table S8.

STARgrid therefore fits the nuclei to the whole profile of counts across the nucleus edge rather than to the count under them alone. It groups the capture locations by their distance from the edge of the nearest nucleus and divides each count by the mean count of the capture locations within 4 µm of it, so that only the local contrast remains. It then moves the nuclei to the shift at which these groups explain the most variation in the counts. This measure, the correlation ratio, comes from the registration of images of different kinds, such as MRI and CT scans, whose intensities are related but not equal (Roche et al. 1998). Masks of the ring alone, or of the nuclei with their ring, each capture only part of this profile (Supplemental Methods). Where the dip could also be used, on the tumour slides, the correlation ratio found the same offset in 303 of 328 slides and regions, among them every whole slide, and an offset one location away in the rest (Supplemental Table S8). Where the dip is weak or absent, the correlation ratio still has a clear maximum (Fig. 4E). We chose the distance of 4 µm for the local mean from distances of 2 to 60 µm; from 20 µm upwards, the correlation ratio also followed the density of the tissue (Supplemental Table S9).

On every slide, the overall registration of Space Ranger version 4, which also uses the counts (10x Genomics 2026d), placed the nuclei where ours does, to the nearest capture location (Fig. 5A). Region by region, however, offsets of up to 4.5 µm remained on the SPATCH slide under our Space Ranger 4.1.0 run (Fig. 5B), whereas STARgrid’s registration, which fits a displacement to the regional offsets, put the nuclei at zero offset in all 22 regions where the offset could be measured (Supplemental Table S8). We also applied the correlation ratio to public slides of 4 other tissues, whose nuclei were placed by Space Ranger 4. It gave a shift on 64 of the 69 slides and regions, each within one capture location of Space Ranger’s registration. These included 11 of the 12 slides and regions without a dip, among them the whole kidney and mouse brain slides (Supplemental Table S10).

Because Space Ranger 4’s registrations also use the counts, judging ours against them is partly circular. As a reference independent of the counts, we used lung tissue whose nuclei Xenium imaged on the same section: 4 cores (Vannan et al. 2025), each registered on its own, and 2 whole sections published by 10x Genomics (Methods). We segmented the nuclei in the H&E image of each in two ways, with Cellpose-SAM and with StarDist. Over the three sets of nuclei, the two from the H&E image and the one imaged by Xenium, the correlation ratio gave a shift on 137 of 152 cores, sections and regions: 109 at the position of the Xenium nuclei, 27 one capture location away and 1 two locations away (Supplemental Fig. S3; Supplemental Table S11). The two segmentations gave comparable results. Where both gave a shift, the two shifts were within one location of each other in 32 of 33 cases, and the shift was at the position of the Xenium nuclei in 25 with Cellpose-SAM and 24 with StarDist. On the whole cores and sections, the correlation ratio brought the StarDist nuclei onto the Xenium nuclei in all 6, and the Cellpose-SAM nuclei in 3, with 2 one location away and the rest without a stable shift. As the two are comparable, STARgrid uses Cellpose-SAM by default, and we used it in all our benchmarks.

### STARgrid adds signal

We first tested the counts that STARgrid adds against the H&E image of the same section, on the colorectal and SPATCH slides and the ovarian dataset (Fig. 6A–C). It is the one reference that shares both the section and the capture grid with the counts. Nuclei were segmented from the image and expanded by a fixed distance to approximate cells, and both masks were placed on the capture grid by STARgrid’s registration (above), which uses neither Space Ranger’s registration nor its counts (Methods). In each 8 µm bin that Space Ranger reports, we split the STARgrid count into the shared counts, the part that both pipelines place there, and the added counts, the remainder.

**Figure 6.**
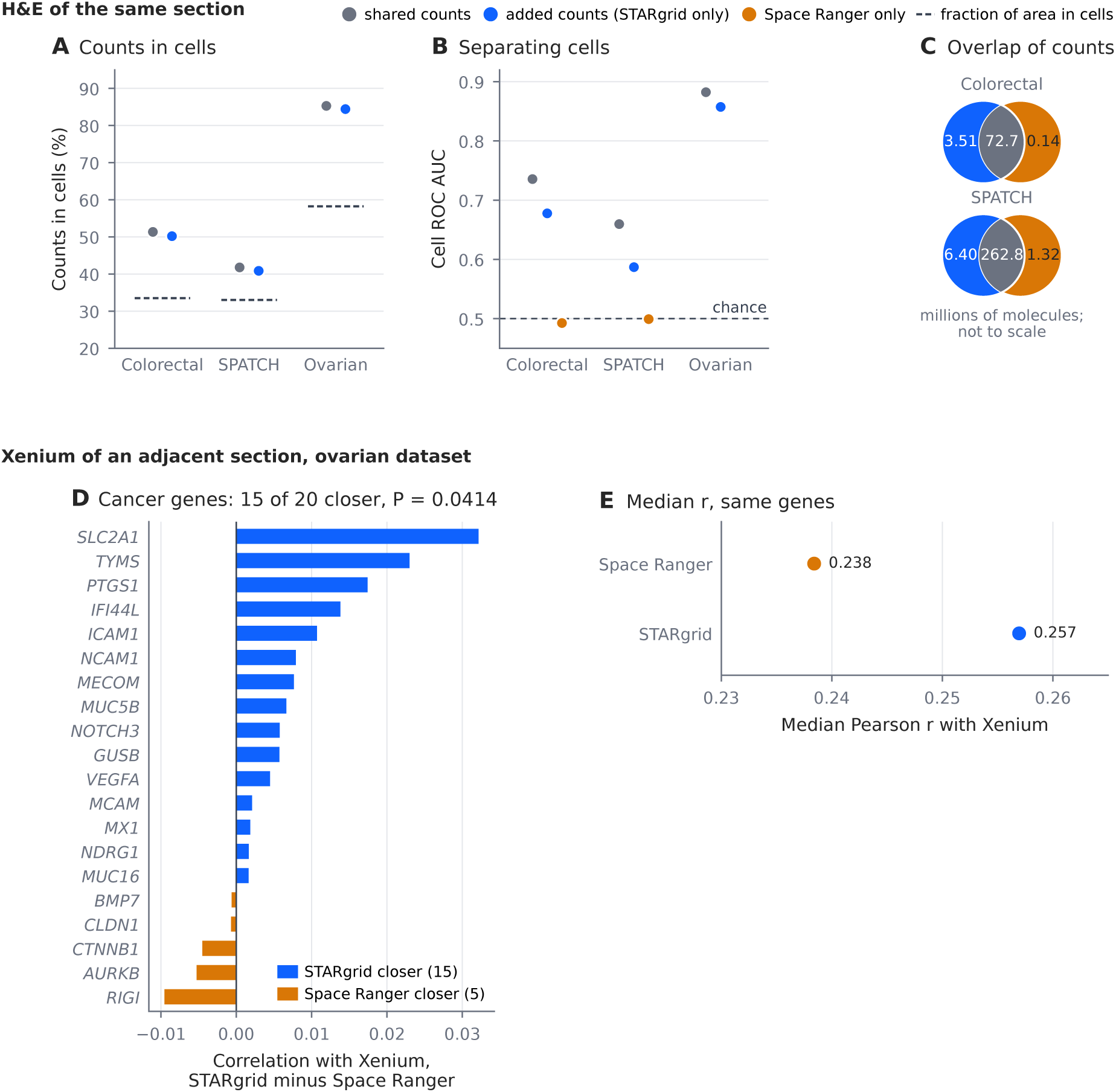
STARgrid adds signal. **A–C**, The H&E image of the same section. Nuclei were segmented from it with Cellpose 4.1.1 (404,441 and 241,920 nuclei on the colorectal and SPATCH slides; the ovarian masks are described in the Supplemental Methods) and expanded to approximate cells, and the masks were placed on the capture grid by STARgrid’s registration (Methods). In each 8 µm bin that Space Ranger reports, the shared counts are the smaller of the two pipelines’ counts; the rest of each pipeline’s count in the bin is counted as that pipeline’s only, and STARgrid’s are the added counts. The comparison is of counts bin by bin, not of matched molecules. **A**, Fraction of the shared and of the added counts that lies on capture locations inside cells; dashed lines, the fraction of locations inside cells, which counts placed at random would give. **B**, How well each set of counts separates locations inside cells from those outside (cell ROC AUC; chance, 0.5). The counts of Space Ranger only, shown for the colorectal and SPATCH slides, lie at chance. **C**, The shared counts and the counts of each pipeline only, on the same two slides, in millions of molecules; the diagrams are not to scale. The counts of Space Ranger only are few, so their ROC AUC in B rests on few counts. These two slides use different Space Ranger comparators (the published raw matrix and the published filtered matrix), and the numbers are not pooled. **D, E**, The ovarian dataset against a Xenium Prime run on an adjacent section of the same block, on 128 µm patches. **D**, For each of the 20 cancer-panel genes with the most Xenium counts, the correlation of STARgrid with Xenium across patches minus that of Space Ranger; exact two-sided sign test. Variants of this comparison are in Supplemental Table S13. **E**, Median correlation with Xenium over the same genes. Absolute correlations are low because the sections are adjacent and registration between them is coarse; only the comparison between the two pipelines is informative.

The fraction of the added counts that lies in cells is nearly that of the shared counts: 1.15 and 0.93 percentage points lower on the colorectal and SPATCH slides, and 0.86 points lower on the ovarian dataset (Fig. 6A; Supplemental Table S12). Both fractions are well above the fraction of the area that cells occupy, which is what random placement would give. Taken alone, the added counts also separate capture locations inside cells from those outside: when a threshold on their density is varied, the area under the ROC curve (ROC AUC) is between 0.587 and 0.857 on these three slides, well above the 0.5 of random placement (Fig. 6B). The values are a little below those of the shared counts, as expected for counts that include the reads hardest to place (Supplemental Table S12).

With the added counts included, STARgrid separates capture locations inside cells from those outside to the same degree as Space Ranger’s counts: the two ROC AUCs differ by 0.000079 on the colorectal slide and 0.000055 on the SPATCH slide. These scores had only a weak dependence on the registration: on the colorectal slide, replacing one step of our registration with Space Ranger’s changed the difference between the pipelines in each H&E score by at most 1.97 *×* 10*^−^*^3^ (Supplemental Fig. S4; Supplemental Table S14).

The added counts also bring the measured expression of cancer genes closer to an independent assay. For the ovarian dataset, a Xenium Prime run was made on an adjacent section of the same block, and we compared the Visium counts with it on 128 µm patches (Fig. 6D,E; Methods). From a cancer gene panel, we took the 20 genes with the most Xenium counts, so that the choice depends on neither pipeline, and for each gene measured how closely each pipeline’s counts rose and fell with Xenium’s from patch to patch (Pearson correlation). STARgrid’s counts followed Xenium’s more closely than Space Ranger’s for 15 of the 20 genes, a significant majority (exact two-sided sign test, *P* = 0.0414; Fig. 6D), with a median correlation of 0.257 against 0.238 (Fig. 6E). STARgrid counts 7.1% more molecules of these genes than Space Ranger and still follows Xenium more closely, so the added molecules fall where Xenium finds these genes, not at random.

## Discussion

STARgrid is, to our knowledge, the first open-source pipeline that performs all four steps for Visium HD, from reads to count matrices registered to the tissue. Because every step is open, anyone can see how reads are placed and counted, change any step, and compare the result with that of other pipelines. The handling of reads whose barcode matches more than one location is an example. STARgrid keeps every matching location until it places the molecule, so the resolution policy that chooses among them can be changed without touching decoding or gene assignment (Methods). In this paper, we focus on HARD, which places each molecule at its most probable location. However, SOFT, which divides a molecule among its possible locations in proportion to their probabilities, may be useful in future work. By keeping the ambiguity in the counts, it could allow downstream positional information to resolve ambiguous barcodes rather than relying on barcode similarity alone.

STARgrid and Space Ranger choose the same location for most reads, and differ in three clearly defined ways. Where the barcodes of two adjacent rows overlap, Space Ranger takes the shorter barcode and STARgrid the one that matches with fewer edits. Space Ranger leaves some reads without a location although their barcodes match one within STARgrid’s limits, and STARgrid places them. When a read matches the coordinate barcodes of more than one capture location equally well, Space Ranger assigns it a location on its own, whereas STARgrid waits until all the reads of the molecule have been grouped and chooses from all of them. With STARgrid being open, these methodologies can be changed and shared with the community to improve the processing.

On a capture grid, assigning transcripts to cells depends first on registering the image to the grid, and this step is not routine. Registration from the images alone was several micrometres off on two of our slides, enough to change which capture locations lie in nuclei, and on the SPATCH slide Space Ranger 4.1.0’s registration left offsets that varied across the slide. We therefore used the counts themselves to make the registration more robust. A profile of counts across the nucleus edge, most often a ring of more molecules just outside each nucleus and fewer under it, was found in every tissue we tested. Because it needs only the counts and the image, it can register published slides as well as new ones. Interesting lines to pursue are whether other patterns in the counts carry more information, whether offsets estimated more locally register better than a displacement fitted to the whole slide, and how the registration performs against nuclei imaged on the same section in tissues other than lung.

STARgrid counts 1.9–7.6% more molecules than Space Ranger on the four slides; on the two slides compared read by read, most of the difference comes from where it places reads (Table 2). Two tests indicate that these extra counts are real signal. In the H&E images, they fall inside cells at nearly the rate of the counts that both pipelines share. Against an independent Xenium measurement of an adjacent section of the ovarian tumour, our counts agreed more closely than Space Ranger’s for 15 of the 20 most abundant genes of a cancer panel. These results are encouraging: a gain of a few percent in molecules, placed in cells, brought the measured expression of cancer genes closer to an independent assay, which suggests that small improvements in how reads become counts can be of biological importance.

## Methods

### Clean-room techniques used to develop STARgrid

Because software derived from Space Ranger could not be redistributed (Introduction), we developed STARgrid as a clean-room implementation, without reading or decompiling any 10x Genomics code. We built it from the documentation that 10x Genomics publishes and from the reference files distributed with Space Ranger: the lists of valid barcodes, the Visium HD slide-design file and the chemistry definition. The documentation gives, for example, the maximum of four edits in a barcode pair (10x Genomics 2026c), and the reference files give the layout of the barcodes on the slide and in Read 1 (Barcode layout). We supplemented these with the output of Space Ranger, published with the datasets or from our own runs: for example, the limit on the edits in each of the two barcodes of a pair, which the documentation does not give, was inferred from it (Spatial barcode decoding). We also used Space Ranger’s output to check our results and to find where the two pipelines differ. Much of the work had already been done for the STAR-Flex and STAR-perturb modules of STAR Suite (Hung et al. 2026), which are also clean-room implementations.

### STARgrid and STAR Suite

Steps 1–3 of STARgrid (Fig. 1) take place in a single run of the STAR Suite program: one command goes from the FASTQ files to the count matrices. Within it, barcode decoding and count assignment were written for STARgrid, and gene assignment uses the methods that STAR Suite already had (Hung et al. 2026). Step 4, registration, was also written for STARgrid and runs as separate programs after this run (Image registration; Software availability).

### Datasets

We used three kinds of public data, listed with their sources and accessions in Supplemental Table S3. Those held in public databases are the SPATCH slide (reads: Genome Sequence Archive for Human, HRA011129; images: BioImage Archive, S-BIAD1900), the Space Ranger output of slide H1-VM2JXXK/A1 (GEO GSE280315) and the lung cores (Visium HD: GEO GSE276934; Xenium: GEO GSE250346); the rest are on the 10x Genomics website. Four complete human Visium HD slides were compared with Space Ranger (Table 1). Xenium data tested the registration and the counts that STARgrid adds: lung tissue measured by Xenium and then by Visium HD on the same section, as tissue cores (Vannan et al. 2025) and as 2 whole sections published by 10x Genomics, and a Xenium Prime 5K run on sections adjacent to the ovarian dataset. Finally, 4 further public Visium HD datasets of other tissues, published by 10x Genomics with Space Ranger’s output and nucleus segmentation, tested the registration; for these, the counts, the capture locations under tissue and the nuclei were taken from the published output.

### Comparison with Space Ranger

Each slide was compared with the Space Ranger output listed in Table 1, from the same reads; STARgrid and our own Space Ranger runs used the probe set or genome reference given there. On the Flex slides, the outputs are compared on the genes that the probe set targets. Space Ranger’s matrices of the two colorectal slides also count molecules on genes whose probes the probe set marks as not included (2.7% and 1.9% of their molecules); these are left out. For the ovarian dataset we ran Space Ranger 4.1.0 ourselves, with introns included, since the STARgrid counts also include introns. The Space Ranger matrices of the colorectal slides and the ovarian dataset are raw: they hold every bin of the capture grid, so the comparison covers the whole slide. The published SPATCH matrix is filtered instead: it holds only the bins that Space Ranger flags as under tissue. On that slide, the comparison is over those bins, and Table 1 also gives the difference from our total over the whole slide. Differences in molecule totals are 100 *×* (STARgrid *−* SR)*/*SR. Agreement is measured by the Pearson correlation of gene totals and of bin totals at each scale, and by the fraction of matrix entries, nonzero in either matrix, that are identical.

### Execution times

Each timed run went from FASTQ files and images to count matrices and the registration of the image, without writing a BAM file or running secondary analysis. Space Ranger does not need nuclei to register the image, so its optional segmentation was turned off. STARgrid used 32 threads, plus a separate rapidgzip process with four threads to decompress each FASTQ file of the pair, and wrote counts under HARD. Its time includes building the barcode lookup tables (Spatial barcode decoding) and both stages of the registration. Because the second stage needs nucleus masks, STARgrid segmented the image with Cellpose-SAM (Pachitariu et al. 2025), a deep-learning model, on a NVIDIA RTX 6000 Ada Generation GPU, while the reads were processed. Times are the medians of three runs of each pipeline, all on the same 32 threads (16 cores) of an AMD Ryzen Threadripper PRO 5965WX processor, with files on an NVMe SSD (Table 1).

### Barcode layout

The Visium HD slide-design file distributed with Space Ranger 4.1.0 lists the 3,350 BC1 sequences with their columns and the 3,350 BC2 sequences with their rows. Read 1 begins with the UMI, which is 9 bases long, followed directly by BC1 and then BC2. With the lengths given in the Results, the barcode of a read is 29 to 31 bases long, and nothing in the read marks where BC1 ends and BC2 begins. We call the pairs of adjacent rows in which the shorter BC2 sequence is the longer one without its first base (Results) overlapping pairs. No pair of BC1 sequences overlaps in this way.

### Spatial barcode decoding

10x Genomics’ documentation states that a barcode pair with no more than four edits (substitutions, deletions or insertions) is mapped to a pair of coordinate barcodes (10x Genomics 2026c), but not how the edits are divided between the two barcodes. In the BAM file that Space Ranger produced for slide H1-VM2JXXK/A1, all but 0.0002% of the locations it recorded can be reached from the read with at most 2 edits in each barcode (Read-by-read comparison with Space Ranger). We therefore allow at most 2 edits in each barcode and four in all.

Because the barcodes vary in length and some overlap, every sequence within 2 edits of each coordinate barcode is generated in advance, in one lookup table for BC1 and one for BC2, built in seconds at the start of each run. For each sequence, a table lists every coordinate barcode that can give it and the number of edits. To decode a read, the decoder looks up each possible BC1 part of Read 1: 13 to 18 bases long, starting at offsets 8 to 12 counted from 0, the range given in the chemistry definition. Without errors, the barcode starts at offset 9, directly after the UMI. For every BC1 found, the decoder then looks up each possible BC2 part that follows it, 12 to 17 bases long. Each pair of barcodes found is a capture location, whose number of edits is the sum for its two barcodes. The decoder keeps every location with the fewest edits, without choosing among them; these are the candidate locations of the read, the best-matching locations of the Results. A read with an N in its barcode is kept, and its barcode is compared directly with every coordinate barcode, with the N counted as one edit; a read with an N in its UMI is dropped.

A read therefore has more than one candidate location only when two or more locations match it with exactly the same number of edits, for example when a missing or extra base can be assigned to either of two neighbouring barcodes (Barcode layout). STARgrid breaks such ties with two kinds of information that the number of edits ignores: the quality score that the sequencer gives each base, since an edit at a base of low quality is more likely to be a sequencing error, and how often each barcode occurs among reads that match a single location with no edits. The tie is broken once for each molecule, from all its reads, rather than for each read (Read cliques and molecule resolution). The Supplemental Methods give the details and the formulas, and where the STAR Suite source computes them.

### Read-by-read comparison with Space Ranger

The comparison of the counts (above) shows where the two pipelines’ counts differ, but not whether they interpret each read in the same way. To check this, we compared them read by read. The count matrices and molecule file that Space Ranger writes record where each molecule was placed, but not which reads it was made from or where each read was placed; only its BAM file records the location of each read. We therefore ran Space Ranger 4.0.1 (reference GRCh38-2024-A) with BAM output on the published FASTQ files of slide H1-VM2JXXK/A1 (721,219,716 read pairs); on the ovarian dataset we used the BAM file of our own Space Ranger run (Comparison with Space Ranger). We kept the reads that Space Ranger assigned to a single gene and gave a UMI, and decoded the Read 1 of each as in a STARgrid run, to obtain its best-matching locations. Each read then falls into one of four cases: Space Ranger’s location is one of these best-matching locations; it is another location; Space Ranger gave the read no location; or STARgrid found no location within its limits but Space Ranger gave one. For every read that Space Ranger placed, we also counted the edits between the read and Space Ranger’s location, on the same part of Read 1 and with no limit on each barcode; these counts set that limit (Spatial barcode decoding). Finally, a STARgrid molecule was matched to a Space Ranger molecule when the two contained exactly the same reads, wherever each pipeline placed them; the matched molecules show how many the two pipelines place at different locations.

### Read cliques and molecule resolution

As in single-cell RNA-seq, reads of the same gene at the same location whose UMIs are identical or differ by one base are counted as one molecule, because a one-base difference between UMIs is more likely to be a sequencing or amplification error than a second molecule (Smith et al. 2017). The two pipelines correct barcodes and group reads in a different order. In the procedure that 10x Genomics documents, the barcode of each read is corrected to a location on its own, and reads are then grouped by location, UMI and gene (10x Genomics 2026c). STARgrid groups the reads before choosing a location: read pairs that have the same gene (Gene assignment, below) and the same raw UMI, and share at least one candidate location, form a read clique, which the Results call a molecule (their numbers on each slide are in Fig. 2A). Each clique is given a location under a resolution policy (next section), from the probability of each of its locations (Supplemental Methods). UMIs that differ by one base are then joined at each location with the directional method of UMI-tools (Smith et al. 2017), and each set of UMIs joined in this way is counted as one molecule. The Supplemental Methods and Hung et al. (2026) give the details.

### Count resolution policies

Because STARgrid keeps every candidate location until the reads have been grouped into cliques, the ties that remain are resolved in a separate step, and one run can resolve them in several ways from the same cliques, without decoding barcodes or assigning genes again. Each way of resolving them is a resolution policy, and the counts under different policies can be compared directly. STARgrid has four. HARD assigns each clique to its most probable location. STRICT keeps only cliques with a single candidate location. SOFT divides each clique among its locations in proportion to their posterior probabilities, so that its counts are expected counts. GATED keeps only cliques whose most probable location passes fixed posterior thresholds, and assigns them as HARD does. The counts under each policy are written as Matrix Market matrices at 2, 8 and 16 µm. Every count reported in the Results uses HARD; the four are compared with Space Ranger in Supplemental Table S2.

### Sources of the molecules STARgrid counts beyond Space Ranger

On slide H1-VM2JXXK/A1 and the ovarian dataset, we divided the difference *D* = *T − M* between STARgrid’s molecule total *T* and the number *M* of molecules that Space Ranger counted into three components: placement, gene assignment and deduplication (Table 2). Each is found by recounting the reads with one factor changed at a time. On slide H1-VM2JXXK/A1, all of these totals count only the genes that the probe set targets, as in the comparison of the counts (Comparison with Space Ranger). The two recounts, *B* and *H*, use the reads that Space Ranger assigned to a single gene and gave a UMI (as in the read-by-read comparison), with Space Ranger’s gene for each read and STARgrid’s correction of their raw UMIs. *B* places these reads at Space Ranger’s locations. *H* places them where STARgrid would, from their candidate locations and read cliques under HARD; it therefore also places the reads that Space Ranger left without a location. *B* differs from *M* only in deduplication, so *B − M* is the deduplication component. *H* differs from *B* only in placement, so *H − B* is the placement component. *T* differs from *H* only in gene assignment, so *T − H* is the gene-assignment component, which includes STARgrid’s selection of reads and its handling of UMIs that have reads for two genes. The three components add up to *D* exactly (Supplemental Table S6).

### Gene assignment

Each read is assigned to a gene, independently of its location, by the methods of STAR Suite (Hung et al. 2026; Dobin et al. 2013; Kaminow et al. 2021), in the same run that decodes its barcode. For RNA-seq quantitation, reads are aligned to the genome, and intronic as well as exonic reads are counted; for Flex quantitation, they are matched to the probes.

### Image registration

STARgrid registers the microscope image to the capture grid in two stages. The first stage aligns the microscope image with the CytAssist image. It finds matching features in the two images with SIFT (Lowe 2004) and fits a similarity transform (shift, rotation and scale) to the matches with RANSAC (random sample consensus), which uses the largest set of matches that agree with one another and ignores the rest (Fischler and Bolles 1981). The CytAssist image is then aligned with the capture grid: the fiducial marks are detected in the image, and a mapping from image to slide coordinates that also corrects for perspective is fitted to their positions in the slide’s Visium layout file, which gives the physical layout of that slide. No Space Ranger alignment, tissue position, mask or count enters this stage.

The second stage fits nucleus masks segmented from the image to the profile of counts across the nucleus edge (Fig. 4), starting from the registration of the first stage. The capture locations are grouped by their distance from the edge of the nearest nucleus, in bands one capture location (2 µm) wide: 3 inside the nuclei and 6 outside them, with the rest of the tissue as one further group. Each count is divided by the mean count of the capture locations under tissue in the 5 *×* 5 square around it (reaching 4 µm on each side of it), which removes changes in the density of the tissue over distances longer than a cell. The groups are then moved, all together, by a candidate offset of a whole number of capture locations in rows and columns, and at each offset the correlation ratio *η*^2^ is computed: the share of the variance of the divided counts that the group means explain (Roche et al. 1998). Every offset within the search is tried, and the masks are moved by the offset with the highest *η*^2^ among the local maxima, the offsets whose *η*^2^ is higher than at each of the 8 offsets one capture location away. If that offset lies on the edge of the search or is not stable when the slide is resampled (Supplemental Methods), the first registration stands and the slide or region is reported as not registered. The offset is also estimated separately in each region of a 5 *×* 5 division of the slide. A displacement that varies linearly with position on the slide is fitted to alternate regions, tested on the regions left out of the fit, and applied.

For the registration, STARgrid segments the nuclei with Cellpose-SAM (Stringer et al. 2021; Pachitariu et al. 2025) (settings in the Supplemental Methods), and the same masks were used to test the added counts (below). To show that the procedure does not depend on the segmentation method, we also registered with nuclei segmented with StarDist (Schmidt et al. 2018), on the ovarian dataset, the two colorectal slides and the lung tissue. On the three tumour slides, the two gave the same whole-slide offsets (Supplemental Table S8). On the lung tissue their shifts were comparable (Results). The counts used were the total HARD count at each 2 µm capture location, summed over genes; on slide H1-VM2JXXK/A1, they were the counts of the Space Ranger run of the read-by-read comparison. Only capture locations that Space Ranger’s output flags as under tissue were used, so the second stage as run depends on this output of Space Ranger. More details are given in the Supplemental Methods.

### Testing the added counts against cells in the H&E image

The cell masks were the nucleus masks of the registration (Image registration), each nucleus expanded by a fixed distance (Supplemental Methods). Scoring was at 8 µm, over every bin in Space Ranger’s matrix, including bins with no counts. In each bin, the shared counts are the smaller of the HARD and Space Ranger counts, and the rest of each pipeline’s count is counted as that pipeline’s only; STARgrid’s are the added counts. Each bin’s count was divided equally among its 2 µm capture locations, and each location was labelled as inside or outside the mask. Two scores were computed. The in-cell fraction is the share of a set of counts on locations inside cells; counts placed at random would give the share of locations that lie inside cells. The ROC AUC is the probability that a location inside the mask has a higher count density than a location outside it, with ties counted as one half; it is 0.5 for random placement whatever the area of the mask.

### Xenium comparison

On the ovarian dataset, STARgrid’s counts and the intron-inclusive Space Ranger 4.1.0 matrix were compared with the Xenium Prime run on an adjacent section. The two sections hold different cells, so they cannot be compared cell by cell; they were compared instead over squares of tissue 128 µm on a side, called patches, laid over the H&E image of the Visium slide. A gene’s expression in a patch was measured separately for each of the three sets of counts: for STARgrid and Space Ranger, as the sum of its counts over the 16 µm bins whose centres lie in the patch, and for Xenium, as the sum over the Xenium cells whose centres lie in it (Supplemental Methods). A patch was evaluated only if it held at least 10 Xenium cells and 16 bins under tissue. Patch expression was log-normalised over the genes present in all matrices.

The genes are the 20 genes of the cancer panel with the most Xenium counts, whose signal is strongest. The panel consists of the custom add-on genes of the Xenium v1 run that 10x Genomics published with the Prime run. For each gene, each pipeline’s patch expression was correlated with Xenium’s across patches (Pearson correlation), and the number of genes whose correlation is higher for STARgrid than for Space Ranger was tested with an exact two-sided sign test. A sensitivity analysis varied the number of genes, the patch size and the correlation measure; every variant is reported, without adjustment for multiple testing (Supplemental Table S13).

### Software availability

STARgrid’s decoding, gene assignment and count assignment are part of STAR Suite (https:// github.com/morphic-bio/STAR-suite; this paper describes release v1.12.2, commit `2ca281c96f3dd2b9d596da1991b397fd05f9f73d`), written in C/C++ and released under the MIT licence for Linux (x86-64). The released binaries need zlib and OpenSSL libcrypto; rapidgzip is optional and decompresses ordinary gzip FASTQ files in parallel. Both stages of the image registration are in https://github.com/BioDepot/stargrid, under the MIT licence (Data access), as the C++ programs that STARgrid runs: `visium_hd_register` for the first stage and `visium_hd_count_register` for the second. The second gives the same results as the Python package stargrid-register in the same repository, which we used for the analyses and which reproduces the shifts reported here. Two scripts run the pipeline on a slide with the settings used here: `scripts/run_visium_hd.sh` in the STAR Suite repository goes from FASTQ files to count matrices, and `scripts/run_registration.sh` in the stargrid repository registers the image. Each has a recipe (`star_visium_hd` and `star_visium_hd_registration`), which lists its settings with their types, defaults and constraints. By agent-ready we mean that an AI agent can set up both without reading the source code or the options of the commands: through the Model Context Protocol server of STAR Suite, it reads the settings, has its values checked and receives the command. The Launchpad shows a person the same recipes as forms (Supplemental Methods). The README files of the two repositories describe how to reproduce a slide of this study from its public data.

## Data access

No new sequencing data were generated in this study. The public datasets analysed, with their accessions and sources, are listed in Methods (Datasets) and Supplemental Table S3. Raw sequencing data, vendor outputs, whole-slide images and licensed third-party software are not redistributed.

The result tables, evaluation scripts and figure-generation code are available at https://github.com/BioDepot/stargrid. The same repository holds, under the MIT licence, the Visium HD tools that are not part of STAR Suite: image registration, nucleus segmentation and the matching of nuclei to the profile of counts around them that the second stage of the registration uses, in C++ and as the Python package stargrid-register in its `register` directory.

STARgrid is released under the MIT licence as part of STAR Suite at https://github.com/morphic-bio/STAR-suite, with precompiled binaries for x86-64 Linux.

## Acknowledgments

We thank 10x Genomics, the SPATCH investigators and the authors of the lung dataset for making their data publicly available. L.H.H. and K.Y.Y. are supported by the National Institutes of Health (NIH) grant U24HG012674. K.Y.Y. is also supported by the Virginia and Prentice Bloedel Endowment at the University of Washington.

## Author contributions

L.H.H. conceived the project, wrote the software, performed the analyses and drafted the manuscript. K.Y.Y. supervised the work and revised the manuscript.

## Supplemental material

**Supplemental Material** (PDF). Supplemental Methods: comparison with Space Ranger, decoding and counting; image registration; evaluation; setting up a run through an AI agent. Supplemental Fig. S1: averages over all nuclei can hide the dip, which appears when the masks are moved. Supplemental Fig. S2: nucleus by nucleus, against a far-shift background. Supplemental Fig. S3: the lung, core by core and section by section, against the nuclei imaged by Xenium. Supplemental Fig. S4: one step of the registration swapped on the colorectal slide. Supplemental Tables: S1, tools that accept raw spatial FASTQ, by the four steps of the pipeline. S2, concordance with Space Ranger for every resolution policy, scale and slide. S3, the datasets used, with their sources. S4, two reads from the read-by-read comparison with Space Ranger. S5, how reads close to more than one capture location are handled. S6, where the molecules STARgrid counts beyond Space Ranger come from, on two slides. S7, nucleus by nucleus: the paired depth of nuclei in place and against a far-shift background. S8, whole-slide and regional offsets under each registration, by the dip and by the correlation ratio. S9, the window of the local mean. S10, the dip and the correlation ratio on four public Visium HD datasets. S11, lung tissue measured by Xenium and then Visium HD on the same section. S12, the in-cell fraction and cell ROC AUC of the shared and the added counts. S13, sensitivity of the top-gene comparison with Xenium to the number of genes, the patch size and the correlation measure. S14, registration sensitivity on the colorectal slide.

