## Supplemental Material for "STARgrid – an open end-to-end pipeline for Visium HD"

### Supplemental Methods. Comparison, decoding and counting

**The barcode lookup.** To find the coordinate barcodes closest to a read in edit distance, the decoder uses two lookup tables, one for BC1 and one for BC2, rather than a single table of barcode pairs. The two tables together cover the 6,700 coordinate barcodes, whereas a table of pairs would have to cover all 11,222,500 capture locations. We confirmed, for every sequence within 2 edits of every coordinate barcode, that the lookup tables give the same answers as calculating the edit distance directly with the Wagner–Fischer algorithm: there were 0 differences.

**The location score.** The location score breaks ties when multiple candidate locations match a read with the same number of edits (main Methods, Spatial barcode decoding). It combines two terms, rather than applying them one after the other: a sequence log-likelihood for each read, which uses the quality of its bases, and a prior on how common each barcode is. The choice is made once for each read clique, from all its reads.

For a read and one of its candidate locations, the sequence log-likelihood is computed from the best alignment of the barcode part of Read 1 that the decoder matched to the location against the location's BC1 and BC2 sequences joined end to end. Each base contributes according to its Phred quality $Q$, through its error probability $e = 10^{-Q/10}$, kept between $10^{-10}$ and 0.75: a base that matches contributes $\log(1-e)$, a substituted base $\log(e/3)$, an extra base in the read $\log(e/4)$ and an N $\log(1/4)$. A base missing from the read has no quality of its own and contributes $\log 10^{-3}$, the error probability at quality 30. When the barcodes can be placed in the read in more than one way that reaches the same location, the placement with the highest log-likelihood is kept. The log-likelihood is the logarithm of the probability of the read's barcode bases if they came from that location, so an edit at a base of low quality, which is more likely to be a sequencing error, lowers it less than an edit at a base of high quality.

The prior uses the reads that decode to a single location with no edits. For a location, $n_c$ is the number of these reads whose column barcode (BC1) is that of the location, and $n_r$ the number whose row barcode (BC2) is that of the location. The prior of the location is $(n_c+1)(n_r+1)$. The counts are multiplied because the two barcodes of a location are treated as independent, so that the prior probability of the location is proportional to that of its column barcode times that of its row barcode; one is added to each count so that a barcode that has no such reads by chance does not get a prior of zero. The counts are taken once, before any tied read is resolved, and are not updated afterwards or carried from one run to another.

For a read clique, the evidence for each location shared by all its reads is the sum of the reads' log-likelihoods for that location plus the logarithm of its prior, $\log(n_c+1)+\log(n_r+1)$. The probability of each location is the exponential of its evidence divided by the sum of the exponentials over the clique's shared locations. In the STAR Suite source, the read log-likelihood is computed by `raw_candidate_alignment_log_likelihood` in `core/legacy/source/SpatialR1Decoder.cpp`, and the clique probabilities by `appendCliquesForGroup` in `core/legacy/source/SpatialGexPipeline.cpp`.

**Read-by-read comparison with Space Ranger.** The comparison used the reads in Space Ranger's BAM file that Space Ranger assigned to a single gene and gave a UMI: the primary alignments with a nonzero `xf` tag, one gene in the `GX` tag, and both a raw (`UR`) and a corrected (`UB`) UMI. Some of these reads belong to molecules that Space Ranger did not count.

Each read was joined by its name to its Read 1 in the FASTQ files, and its barcode was decoded with the lookup tables above, over starts 8 to 12 and barcode lengths 25 to 35. The location that Space Ranger recorded for a read is its `CB` tag. The edits between the read and that location were counted for every start and for every point at which the barcode can be split into its two parts. Of the 639,672,962 locations that Space Ranger recorded, all but 1,483 (0.0002%) can be reached from the read, for some start and split, with at most 2 edits in each barcode; this is the basis of the limit of 2 edits per barcode that the decoder uses.

The molecules that Space Ranger counted are those with a representative read marked in the `xf` tag; they are the molecules of its `molecule_info.h5` file.

**Read cliques.** Read pairs that have a gene and at least one candidate location are first grouped by gene and raw UMI. Each group is then divided into read cliques by a deterministic procedure: a read pair joins an existing clique only if one of its candidate locations is shared by every read pair already in that clique, and otherwise it starts a new clique. Read pairs linked only through a chain of shared locations, for example A sharing a location with B and B with C, but A none with C, therefore end up in separate cliques.

Cliques are formed from the raw UMI, before any correction of UMIs, because correcting UMIs first could merge the reads of two different molecules whose UMIs happen to differ by one base. A sequencing error in a raw UMI, by contrast, only splits the reads of one molecule into two cliques, which the later correction of UMIs can join again.

**Sources of the molecules STARgrid counts beyond Space Ranger.** On slide H1-VM2JXXK/A1, $M$ is the number of molecules that our Space Ranger 4.0.1 run of the read-by-read comparison counted on the genes that its probe set targets, and $B$ and $H$ count the same genes. $T$ is the total of a STARgrid run with the settings used for the colorectal slide and the same probe set; it needs no such restriction, because every gene that STARgrid counts is targeted by the probe set. On the ovarian dataset, $M$ is the number counted by the Space Ranger 4.1.0 run with introns included that the other comparisons use, and $T$ is the total of the STARgrid run reported in the paper. The recounts $B$ and $H$ correct UMIs in the same way as the matrices, without moving UMIs from one gene to another.

### Supplemental Methods. Image registration

Shifts are given as (rows, columns) of capture locations, with rows running down the slide, and measure the move from the nucleus masks to the counts. Supplemental Figs. S1–S3 and Supplemental Tables S7–S11 hold the values behind these methods.

**Registration of the microscope image to the capture grid.** The main Methods describe the two steps of the first stage, which uses the images only. Before features are matched, the microscope image is reduced to about the pixel size of the CytAssist image and converted to a haematoxylin-like intensity. The SIFT features of the two images are matched in every orientation of the microscope image, including reflections, and a match is kept only if it is clearly better than the next best match (a ratio test on the feature descriptors). Once the similarity transform has been fitted to the matches, it is not refined further. The mapping that aligns the CytAssist image with the capture grid is accepted only if a minimum number of fiducial marks fit it.

**The correlation ratio.** Each capture location is assigned to a band by its Euclidean distance from the nucleus edge: under a nucleus, at most one location from the edge, one to two locations, or more than two; outside the nuclei, within $k-1$ to $k$ locations of a nucleus, for $k = 1, \dots, 6$; the other capture locations under tissue form one more group. Moving the nucleus masks moves the bands with them. Each count is divided by the mean count of the capture locations under tissue in the $5 \times 5$ square centred on it. At each shift, over the capture locations under tissue of the slide or region, $\eta^2 = \sum_g n_g (m_g - m)^2 \big/ \sum_i (v_i - m)^2$, where $v_i$ is the divided count of location $i$, $m$ the mean of the $v_i$, and $n_g$ and $m_g$ the number and the mean of the $v_i$ in group $g$; a location counts under the band that the shift brings onto it. $\eta^2$ is zero when every group has the same mean and one when the divided counts are constant within each group. It does not require any group to hold more or fewer counts than another.

**The search.** The search covered shifts of up to 12 locations in each direction in every analysis: whole slides, regions and lung cores. The shift found is the highest local maximum of $\eta^2$, higher than at each of the 8 neighbouring shifts. A whole slide, or a region, was usable if this shift lay inside the search and was stable: when the slide was resampled 200 times in spatial blocks, the shift found lay within one location of the same shift in at least 95% of the resamples. The dip was found and tested in the same way, as the deepest local minimum of the mean count under the nuclei.

**From the dip to the correlation ratio.** We first registered with the dip alone. On the two whole lung sections it lay at the position of the Xenium nuclei but was not stable, and on the kidney and mouse brain slides there was none. We then moved a mask of the ring, the capture locations within 3 locations (6 µm) of a nucleus that are not under one, and took its highest peak; this brought the nuclei onto the Xenium nuclei on both whole sections, but on the kidney and mouse brain slides the peak of the ring moved with its width. On those slides the nuclei and their ring together, the disk, had a peak within one location of Space Ranger's registration. The dip, the ring and the disk each weigh part of the profile of counts across the nucleus edge, with a choice of which part should hold more counts and which fewer. The correlation ratio uses the whole profile without that choice. Dividing each count by its local mean removes the changes in the density of the tissue over distances longer than a cell; without it, fewer slides and regions gave a shift, and on some tumour regions the shift lay two or more locations from a usable dip (Supplemental Table S9, first row).

**The window of the local mean.** We computed the correlation ratio on every slide, region and lung core above with square windows of half-width 1, 2, 3, 4, 5, 7, 10, 15, 20 and 30 locations (2 to 60 µm) and without the division (Supplemental Table S9). We use 2 locations (4 µm), the smallest of the windows that gave the most shifts at the position of the Xenium nuclei on the lung tissue. The smallest window, which holds few counts, gave fewer shifts; from 20 µm upwards, the shift of more and more tumour slides and regions lay two or more locations from a usable dip, up to 22 at the largest window, because a window that spans many cells passes changes in the density of the tissue on to the correlation ratio.

**Regional offsets and the fitted displacement.** Regional offsets were measured in each region of a $5 \times 5$ division of the slide. A region was usable if it held at least 30,000 capture locations under nuclei and its shift was usable as defined above. A displacement that varies linearly with position on the slide was fitted to the offsets of alternate regions and tested on the regions left out of the fit. On the SPATCH slide, it predicted all of them to within one location on each axis, and the masks were moved by this displacement.

**Space Ranger's registrations.** To place our masks by Space Ranger's registration, we fitted a projective transform to the positions in the microscope image that Space Ranger's output gives for its 2 µm capture locations or, in the output deposited with slide H1-VM2JXXK/A1, for the centres of its 8 µm bins. We used the outputs deposited with the ovarian dataset (Space Ranger 4.0.1), the colorectal slide (4.1.0) and slide H1-VM2JXXK/A1 (3.0.0, in GEO), and our own runs of Space Ranger on a subsample of 100,000 reads, each given the same image as the deposited output: versions 3.0.0 and 4.1.0 on the SPATCH slide, with the same supplied manual alignment, and version 4.1.0 on slide H1-VM2JXXK/A1, with a manual alignment derived from the GEO deposit. The second stage used only the capture locations that Space Ranger's output flags as under tissue; the published output of the SPATCH slide has no flags at 2 µm, so for that slide the flags came from our runs on the subsample.

**Lung tissue measured by Xenium and by Visium HD on the same section.** The public lung dataset of Vannan et al. (cited in the main text) was measured by Xenium and then by Visium HD on the same section, in 4 tissue cores of slide H1-K3WKZHN/D1, and so were the 2 whole sections published by 10x Genomics (Supplemental Table S3). Because both assays measured the same section, the nuclei imaged by Xenium show where the nuclei lie on the capture grid without using the counts around nuclei. We placed the Xenium nucleus boundaries on the capture grid with a transform fitted to the density of each gene's molecules in the two assays (on the second section, the genes with the most molecules), using neither nuclei, nor the H&E image, nor the pattern of total counts around nuclei. On the whole sections, which extend beyond the capture area, only the capture locations inside the area imaged by Xenium and covered by the H&E image were used.

The nuclei tested were segmented from the dataset's H&E image with Cellpose-SAM, with the settings used on the colorectal slide (Supplemental Methods, Evaluation), and, as a control, with StarDist; both were placed by the dataset's own registration, made with Space Ranger 3.0.0 (by hand for the cores and the second section). The shift that brings them onto the Xenium nuclei was measured for each segmentation by cross-correlating the two sets of masks, without counts. The test is whether the shift found for the H&E nuclei in the counts deposited with the dataset lies at this shift, and whether the shift found for the Xenium nuclei is zero (Supplemental Table S11). Each core was registered and tested on its own, with only its own capture locations, because the cores lie at different offsets from the dataset's registration. Regions were those of a $3 \times 3$ division of each core and a $5 \times 5$ division of each whole section, each holding at least 30,000 capture locations under nuclei.

**The dip, nucleus by nucleus.** To test the dip nucleus by nucleus, we computed a paired depth for each nucleus of at least 4 capture locations: one minus the mean count on its capture locations under tissue divided by the mean count on the same locations moved to each of the 8 neighbouring shifts, or to the 16 next shifts out. Its median over nuclei is reported, with intervals from 200 draws of spatial blocks of the slide. Because this statistic is above zero even for masks moved far from their nuclei, the same comparison was made with each nucleus first moved 25 locations (50 µm on each axis) along each of the 4 diagonals, and this background was subtracted (Supplemental Fig. S2; Supplemental Table S7). The tumour nuclei tested are the nucleus maps deposited with the ovarian dataset and the two colorectal slides, and our Cellpose masks of the ovarian dataset moved by $(-3, -3)$; the lung nuclei are those imaged by Xenium.

### Supplemental Methods. Evaluation

**Testing the added counts against cells in the H&E image.** On the colorectal and SPATCH slides, nuclei were segmented with Cellpose 4.1.1 (model `cpsam`, Ruifrok-style hematoxylin optical density channel; flow threshold 0.4, cell-probability threshold 0, minimum size 15 pixels, maximum size fraction 0.4). Cells were formed by expanding each nucleus by 2 µm, with each pixel assigned to the nearest nucleus. The image was segmented in tiles of 2,048 pixels overlapping by 128 pixels, with a fixed rule deciding which tile owns each overlap.

The ovarian masks were made with Cellpose 4.1.1 for an earlier analysis of that slide, with a flow threshold of 0.8 and an 8 µm expansion of each nucleus. They were placed with our registration from that analysis before being moved by the second stage of the registration.

After the masks of the SPATCH slide had been moved by the second stage, 882 of the 8 µm bins reported by both pipelines, at the edge of the image, had no mask. These bins hold about 0.07% of each set of counts and were left out of the H&E scores of both pipelines.

**The H&E scores under two registrations.** To test whether the H&E scores depend on the registration, the colorectal slide was also scored against the cell segmentation that 10x Genomics distributes with the dataset. The segmentation was placed on the capture grid twice: once with Space Ranger's registration of the microscope image to the CytAssist image, and once with ours. Everything else, including the placement of the CytAssist image on the capture grid, was the same in the two scorings (Supplemental Fig. S4; Supplemental Table S14).

**Xenium comparison.** The 16 µm bins were placed on the H&E image of the Visium slide with their positions in the Space Ranger output published with the dataset; the counts of both pipelines are in the same bins. Each Xenium cell centre was taken from Xenium coordinates to the H&E image of the Xenium section with the image alignment published with the Xenium data, and from there to the H&E image of the Visium slide with an affine transform fitted to the haematoxylin intensity of the two images, without expression data. Because the sections are adjacent, this registration is coarse, and it does not match cells between the sections. The variants of the comparison are defined in Supplemental Table S13.

#### Supplemental Methods. Setting up a run through an AI agent

STAR Suite offers its recipes to AI agents through a Model Context Protocol server, and to people through the Launchpad, a form in a web browser. A recipe is a file that lists the settings of one kind of run, with the type, the default and the constraints of each, and both routes give the same command. STARgrid has two recipes. That of its STAR Suite run, from FASTQ files to count matrices, is `star_visium_hd`. Its settings are the method of quantitation, the folder of FASTQ files, the slide-design file or a folder of barcode inputs written from it, the output folder, the genome index or the Flex reference, and the settings of the run, such as the number of threads. That of the registration is `star_visium_hd_registration`. Its settings are the microscope image, the CytAssist image, the layout file and the capture area of the slide, the folder of count matrices written by the run, the output folder, and the settings of the segmentation of the nuclei.

An agent needs neither the source code nor the command-line options. It asks the server for the recipes, reads the settings of one, has its values checked and receives the command. Values that break a constraint are refused with the reason, so the agent can correct them before anything is run. The calls are shown below for the run and for the registration of the ovarian dataset, with the paths replaced by names. In the first check of the run, the slide-design file was left out and the number of threads was zero; in that of the registration, the image file did not exist and the number of threads was zero. The command that the agent receives for the run starts STAR Suite with the same options as our own run of `scripts/run_visium_hd.sh` on this dataset, which reproduced the count matrices of the manuscript. The command that it receives for the registration gave the same registered positions as our own run of `scripts/run_registration.sh` in the stargrid repository.

```
list_workflows
  -> star_visium_hd: STARgrid Visium HD (FASTQ to matrices)

get_workflow_parameter_schema  star_visium_hd
  -> mode, fastq_dir, sample, slide, contract, out_dir, genome_dir, flex_reference,
     probe_set, gtf, threads, reads, memory_fraction, star, dry_run
     required: mode, fastq_dir, out_dir

validate_workflow_parameters  mode = gex, fastq_dir = <FASTQ folder>,
     out_dir = <output folder>, genome_dir = <genome index>, threads = 0
  -> valid: False
     At least one parameter must be provided from: slide, contract. Give the
     slide-design file, or a folder that already holds the barcode inputs.
     Parameter 'threads' must be positive, got: 0

validate_workflow_parameters  the same, with slide = <slide-design file>, threads = 12
  -> valid: True

render_workflow_command  the same values
  -> scripts/run_visium_hd.sh --mode gex --fastq-dir <FASTQ folder>
     --slide <slide-design file> --out-dir <output folder>
     --genome-dir <genome index> --threads 12

get_workflow_parameter_schema  star_visium_hd_registration
  -> stargrid, he, cytassist, layout, area, counts, out_dir, segmentation, pixel_size_um,
     cellpose_model, python, container, cpu, threads, bin_dir, dry_run
     required: stargrid, he, cytassist, layout, area, counts, out_dir

validate_workflow_parameters  stargrid = <stargrid repository>, he = <H&E image>.missing,
```

```
    cytassist = <CytAssist image>, layout = <layout file>, area = D1,
    counts = <count matrices>, out_dir = <output folder>, container = <Cellpose image>,
    threads = 0, bin_dir = <registration programs>
  -> valid: False
    Parameter 'threads' must be >= 1, got: 0
    Parameter 'he' path does not exist: <H&E image>.missing

validate_workflow_parameters  the same, with he = <H&E image>, threads = 12
  -> valid: True

render_workflow_command  the same values
  -> scripts/run_visium_hd_registration.sh --stargrid <stargrid repository>
    --he <H&E image> --cytassist <CytAssist image> --layout <layout file> --area D1
    --counts <count matrices> --out-dir <output folder> --container <Cellpose image>
    --threads 12 --bin-dir <registration programs>
```

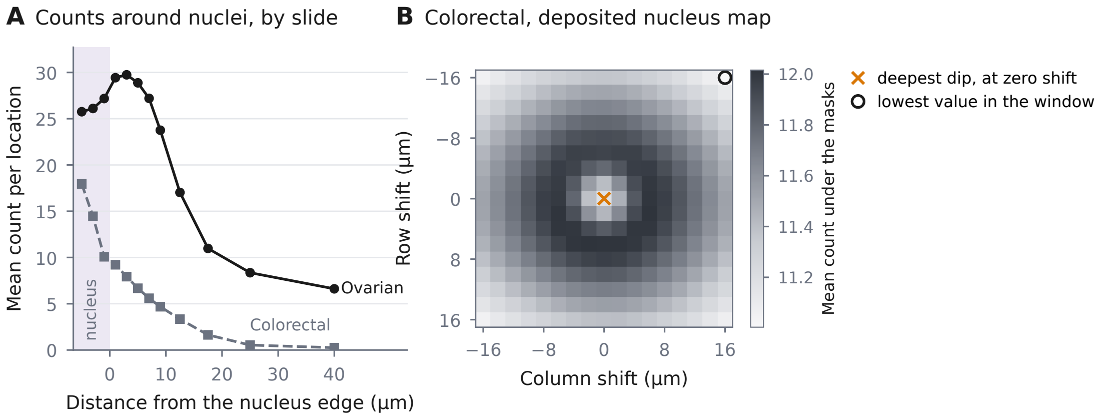

**Supplemental Fig. S1. Averages over all nuclei can hide the dip, which appears when the masks are moved. A**, Mean count per capture location against distance from the edge of the nearest nucleus, in the nucleus maps deposited with the ovarian dataset and with the colorectal slide; negative distances are inside nuclei (shaded). On the ovarian dataset the count rises from inside the nuclei to a peak just outside them; on the colorectal slide it is highest deep inside nuclei and falls outwards, although the colorectal nuclei show the dip when the masks are compared with themselves moved (Supplemental Fig. S2). The open-ended outermost distance bin is not drawn, and the innermost bin, open-ended towards the centre of the nucleus, is drawn one bin width inside the next. **B**, Mean count under the nucleus map deposited with the colorectal slide at each shift up to 16 µm on each axis; rows run down the slide. The count at zero shift (cross) is lower than at every neighbouring shift, a dip, but the lowest value over the shifts shown lies at a corner of the window (circle). The dip is therefore found against its neighbouring shifts (Supplemental Methods), not as the lowest value of the search. Both panels are post hoc and exploratory.

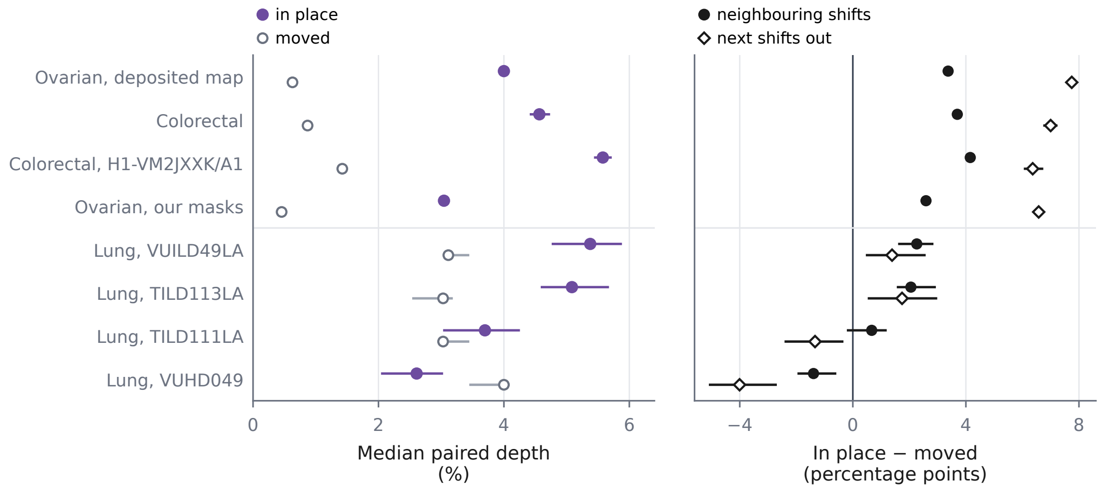

**Supplemental Fig. S2. The dip nucleus by nucleus, against a far-shift background.** Paired depth of each nucleus, as defined in the Supplemental Methods, summarised as the median over nuclei. Left, against the 8 neighbouring shifts: nuclei in place (filled circles) and the same nuclei moved 25 locations (50 µm on each axis) along each of the 4 diagonals (open circles), the background. Right, in place minus background, in percentage points, against the neighbouring shifts (filled circles) and the 16 next shifts out (open diamonds). Bars, 95% intervals from 200 draws of spatial blocks of the slide; most are narrower than the markers. The rows "Ovarian, deposited map", "Colorectal" and "Colorectal, H1-VM2JXXK/A1" use the nucleus maps deposited with those datasets; "Ovarian, our masks" uses our Cellpose masks, placed by our registration and moved by its second stage; the lung rows use the nuclei imaged by Xenium on the same section as the Visium HD run (Supplemental Fig. S3). Values are in Supplemental Table S7.

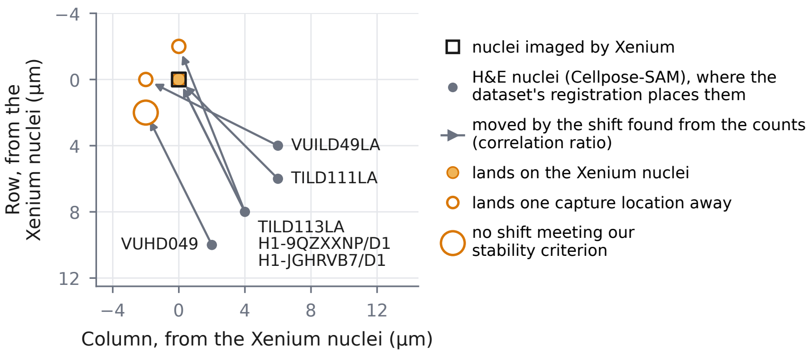

**Supplemental Fig. S3. The lung, core by core and section by section: the registration from the counts against the nuclei imaged by Xenium.** Lung tissue measured by Xenium and then Visium HD on the same section: 4 cores of slide H1-K3WKZHN/D1 (Vannan et al., cited in the main text), each registered on its own, and the whole sections of slides H1-9QZXXNP/D1 and H1-JGHRVB7/D1 published by 10x Genomics. Square, the nuclei imaged by Xenium. Grey dots, where the dataset's own registration, made with Space Ranger 3.0.0 (by hand for the cores and for slide H1-JGHRVB7/D1), places the nuclei segmented with Cellpose-SAM from the H&E image, relative to the Xenium nuclei, measured from the two sets of masks without counts. Arrows, the shift that the correlation ratio finds for these nuclei in the counts deposited with the dataset. Filled circle, the shift lands on the Xenium nuclei; small open circle, one capture location away; large open circle, no shift meets our stability criterion (Supplemental Methods). Rows run down the slide. The values, with the StarDist nuclei as the control, are in Supplemental Table S11.

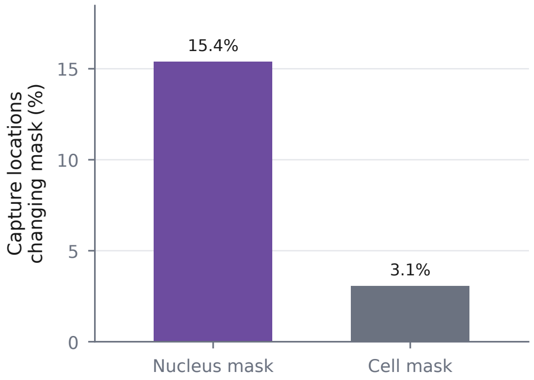

**Supplemental Fig. S4. One step of the registration swapped on the colorectal slide.** Share of capture locations that move into or out of a nucleus mask, and into or out of a cell mask, on the colorectal slide when our registration of the microscope image to the CytAssist image is replaced by Space Ranger's, with the placement of the CytAssist image on the capture grid unchanged; capture locations move a median of 2.70 µm. The difference between the pipelines in each H&E score changes by at most $1.97 \times 10^{-3}$ (Supplemental Table S14).

**Supplemental Table S1. Tools that accept raw spatial FASTQ, by the four steps of the pipeline.** Whether each reviewed tool documents Visium HD barcode decoding, gene assignment for both RNA-seq and Flex quantitation, binned counts at 2, 8 and 16 µm, and registration of the tissue image to the capture grid, with its licence and whether its counts depend on vendor output. "Not documented" describes the reviewed version only. Vendor pipelines are described from their licences, official documentation and observed outputs.

| Tool | HD barcode decoding | RNA-seq + Flex gene assignment | 2/8/16 µm counts | Image registration | Open licence | Vendor-count independent |
|---|---|---|---|---|---|---|
| STARgrid (this work) | yes | yes | yes | yes, from the counts under nuclei | MIT | yes |
| Space Ranger 4.1 | yes | yes | yes | yes, refined against the counts | no | — |
| INTERSTELLAR (Kijima et al. 2023) | not documented | not documented | not documented | not documented | yes | no |
| spacemake (Sztanka-Toth et al. 2022) | not documented | RNA-seq only | not documented | H&E to an image of the counts | yes | yes |
| stPipe 1.2.0 (Xu et al. 2025) | not documented | yes | not documented | not documented | yes | yes |
| RAD (Vaidya et al. 2026) | long reads only | no | no | no | yes | — |

**Supplemental Table S2. Concordance with Space Ranger for every resolution policy, scale and slide.** Mass ratio is the STARgrid molecule total divided by the Space Ranger total. Exact entries are the fraction of entries nonzero in either matrix that have identical counts. Correlations are Pearson *r*. SPATCH uses the bins in Space Ranger's published filtered matrix at 8 µm.

| Slide | Bin (µm) | Policy | Mass ratio | Exact entries | Gene-total *r* | Bin-total *r* |
|---|---|---|---|---|---|---|
| Colorectal | 2 | STRICT | 0.8815 | 0.8109 | 0.999998 | 0.98046 |
| Colorectal | 2 | SOFT | 1.0454 | 0.6589 | 0.999999 | 0.98373 |
| Colorectal | 2 | HARD | 1.0462 | 0.8456 | 0.999999 | 0.98430 |
| Colorectal | 2 | GATED | 0.9515 | 0.8197 | 0.999999 | 0.97900 |
| Colorectal | 8 | STRICT | 0.8815 | 0.8091 | 0.999998 | 0.99848 |
| Colorectal | 8 | SOFT | 1.0454 | 0.8015 | 0.999999 | 0.99910 |
| Colorectal | 8 | HARD | 1.0462 | 0.9045 | 0.999999 | 0.99926 |
| Colorectal | 8 | GATED | 0.9515 | 0.8548 | 0.999999 | 0.99877 |
| Colorectal | 16 | STRICT | 0.8815 | 0.7993 | 0.999998 | 0.99950 |
| Colorectal | 16 | SOFT | 1.0454 | 0.8484 | 0.999999 | 0.99975 |
| Colorectal | 16 | HARD | 1.0462 | 0.9148 | 0.999999 | 0.99980 |
| Colorectal | 16 | GATED | 0.9515 | 0.8551 | 0.999999 | 0.99964 |
| H1-VM2JXXK/A1 | 2 | STRICT | 0.7953 | 0.7462 | 0.9999981 | 0.97975 |
| H1-VM2JXXK/A1 | 2 | SOFT | 1.0262 | 0.5581 | 0.9999997 | 0.98744 |
| H1-VM2JXXK/A1 | 2 | HARD | 1.0263 | 0.8850 | 0.9999997 | 0.98916 |
| H1-VM2JXXK/A1 | 2 | GATED | 0.8783 | 0.8063 | 0.9999959 | 0.98431 |
| H1-VM2JXXK/A1 | 8 | STRICT | 0.7953 | 0.7359 | 0.9999981 | 0.99802 |
| H1-VM2JXXK/A1 | 8 | SOFT | 1.0262 | 0.7322 | 0.9999997 | 0.99901 |
| H1-VM2JXXK/A1 | 8 | HARD | 1.0263 | 0.9203 | 0.9999997 | 0.99935 |
| H1-VM2JXXK/A1 | 8 | GATED | 0.8783 | 0.8103 | 0.9999959 | 0.99853 |
| H1-VM2JXXK/A1 | 16 | STRICT | 0.7953 | 0.7072 | 0.9999981 | 0.99924 |
| H1-VM2JXXK/A1 | 16 | SOFT | 1.0262 | 0.7914 | 0.9999997 | 0.99971 |
| H1-VM2JXXK/A1 | 16 | HARD | 1.0263 | 0.9237 | 0.9999997 | 0.99981 |
| H1-VM2JXXK/A1 | 16 | GATED | 0.8783 | 0.7905 | 0.9999959 | 0.99948 |
| Ovarian | 2 | STRICT | 0.8760 | 0.7633 | 0.998937 | 0.98186 |
| Ovarian | 2 | SOFT | 1.0735 | 0.5881 | 0.998913 | 0.98429 |
| Ovarian | 2 | HARD | 1.0764 | 0.8028 | 0.998925 | 0.98608 |
| Ovarian | 2 | GATED | 0.9577 | 0.7748 | 0.998949 | 0.97943 |
| Ovarian | 8 | STRICT | 0.8760 | 0.7427 | 0.998937 | 0.99854 |
| Ovarian | 8 | SOFT | 1.0735 | 0.7212 | 0.998913 | 0.99910 |
| Ovarian | 8 | HARD | 1.0764 | 0.8543 | 0.998925 | 0.99936 |
| Ovarian | 8 | GATED | 0.9577 | 0.7975 | 0.998949 | 0.99882 |
| Ovarian | 16 | STRICT | 0.8760 | 0.7214 | 0.998937 | 0.99943 |
| Ovarian | 16 | SOFT | 1.0735 | 0.7660 | 0.998913 | 0.99971 |
| Ovarian | 16 | HARD | 1.0764 | 0.8564 | 0.998925 | 0.99979 |
| Ovarian | 16 | GATED | 0.9577 | 0.7878 | 0.998949 | 0.99961 |
| SPATCH | 8 | STRICT | 0.8054 | — | 0.999995 | 0.99663 |
| SPATCH | 8 | SOFT | 1.0229 | — | 0.999997 | 0.99841 |
| SPATCH | 8 | HARD | 1.0232 | — | 0.999997 | 0.99877 |
| SPATCH | 8 | GATED | 0.8876 | — | 0.999996 | 0.99756 |

**Supplemental Table S3. Datasets.** Every dataset used, with its source and what it was used for. The four Visium HD slides compared with Space Ranger are described further in Table 1 of the main text. A dash marks a slide number not given with the dataset.

| Dataset | Slide | Assay | Source | Used for |
|---|---|---|---|---|
| *Visium HD slides compared with Space Ranger* | | | | |
| Colorectal cancer, FFPE | H1-GMHFWPH/D1 | Visium HD, Flex | 10x Genomics: Visium HD Flex human colorectal cancer | comparison, timing, H&E tests, registration |
| SPATCH ovarian cancer, FFPE (Ren et al. 2025) | H1-WP2FTZD/D1 | Visium HD, Flex | SPATCH resource: reads in the Genome Sequence Archive for Human (HRA011129), images in the BioImage Archive (S-BIAD1900), filtered count matrix from `https://spatch.pku-genomics.org` | comparison, timing, H&E tests, registration |
| Ovarian cancer, fresh frozen | H1-YQJBZ7X/D1 | Visium HD 3′ | 10x Genomics: Visium HD 3′ human ovarian cancer, fresh frozen (minimum-depth library) | comparison, timing, H&E and Xenium tests, read-by-read comparison, registration |
| Colorectal cancer, FFPE | H1-VM2JXXK/A1 | Visium HD, Flex | 10x Genomics: Visium HD human colorectal cancer; Space Ranger output also in GEO (GSE280315, sample GSM8594568) | comparison, read-by-read comparison, registration |
| *Xenium* | | | | |
| Ovarian cancer, fresh frozen, sections adjacent to the Visium HD slide | — | Xenium Prime 5K (200,900 cells) | 10x Genomics: Cross-Platform Comparison: FF Human Ovarian Cancer with Xenium v1 and Xenium Prime 5K (resource `Xenium_Prime_Human_Ovary_Cancer_FF`; CC BY 4.0) | Xenium tests of the added counts |
| Lung tissue cores (Vannan et al. 2025) | H1-K3WKZHN/D1 | Xenium, then Visium HD on the same section | GEO: Visium HD with Space Ranger's counts and registration in GSE276934 (sample GSM8509590); Xenium in GSE250346 | registration test |
| Lung adenocarcinoma, whole section, Experiment 1 | H1-9QZXXNP/D1 | Xenium v1, then Visium HD on the same section | 10x Genomics, with Space Ranger 3.0.0's counts and registration: Visium HD post-Xenium human lung cancer and Xenium human lung cancer (CC BY 4.0) | registration test |
| Lung adenocarcinoma, whole section, Experiment 2 | H1-JGHRVB7/D1 | Xenium Prime 5K, then Visium HD on the same section | as Experiment 1; registered by hand | registration test |
| *Other tissues, published with Space Ranger 4.0.1's output and nucleus segmentation (CC BY 4.0)* | | | | |
| Human lymph node, FFPE | — | Visium HD, Flex | 10x Genomics: human lymph node | registration in other tissues |
| Human prostate cancer, FFPE | — | Visium HD, Flex | 10x Genomics: human prostate cancer | registration in other tissues |
| Human kidney, FFPE | — | Visium HD, Flex | 10x Genomics: human kidney | registration in other tissues |
| Mouse brain | — | Visium HD, Flex | 10x Genomics: mouse brain | registration in other tissues |

**Supplemental Table S4. Two reads from the read-by-read comparison with Space Ranger.** Each read's barcode in Read 1 is shown split between BC1 and BC2 (|); rows index BC2 and columns BC1 on the 2 µm grid. In both reads the best-matching location needs one edit: the BC1 segment is one base shorter than its 15-base oligonucleotide, and the BC2 segment matches a 15-base oligonucleotide exactly. Space Ranger's location keeps the same BC1 and uses the 14-base BC2 oligonucleotide of the adjacent row, which lacks the underlined first base, so the read needs an inserted base: two edits. The two locations are 2 µm apart. In read 1 they share an 8 µm bin; in read 2 they lie on either side of a 16 µm bin boundary. In both reads the two BC2 oligonucleotides overlap, as they do for 182 pairs of adjacent rows in the design. The reads are the first in the output of the read-by-read comparison, among those with a single best-matching location, that meet these criteria.

| Location | Row | Column | BC1 (edits) | BC2 (edits) | Total edits |
|---|---|---|---|---|---|
| Read 1, raw barcode `GTCTGCTCCTTTCG|TATTGGCGCATCAAG` | | | | |
| best match | 1183 | 2930 | `GTCTGCTCCTTATCG` (1) | `TATTGGCGCATCAAG` (0) | 1 |
| Space Ranger | 1182 | 2930 | `GTCTGCTCCTTATCG` (1) | `ATTGGCGCATCAAG` (1) | 2 |
| Read 2, raw barcode `GTCAGGAATCCCAG|CATGCAATCAAATAG` | | | | |
| best match | 727 | 2736 | `GTCAGGAATGCCCAG` (1) | `CATGCAATCAAATAG` (0) | 1 |
| Space Ranger | 728 | 2736 | `GTCAGGAATGCCCAG` (1) | `ATGCAATCAAATAG` (1) | 2 |

**Supplemental Table S5. How reads close to more than one capture location are handled.** Space Ranger's entries describe its slide-design reference file distributed with version 4.1.0, its published documentation, and its recorded output from version 4.0.1 on slide H1-VM2JXXK/A1. Where the documentation gives no rule, the entry describes the observed output; the reason for it cannot be checked without the implementation. STARgrid entries describe its stated rule (Methods).

| | Space Ranger | STARgrid |
|---|---|---|
| Barcode oligonucleotides | Slide-design file: BC1 15 or 16 bases, BC2 14 or 15, alternating between adjacent locations | |
| Correction tolerance | "Up to four edits are permissible to correct a barcode to the inclusion list"; Visium HD "follows a similar algorithm" to Visium v1 and v2, where a posterior from base qualities decides whether to correct. Recorded output: all but 0.0002% of emitted locations lie within 2 edits per half | Up to 2 edits per half, within the documented total |
| Output per read | One location (`CB`) or none | Every location with the fewest edits in total, summed over BC1 and BC2; locations with more edits are dropped, and the choice among the tied locations is made per molecule |
| Rule for a read that matches two locations | Not stated in the documentation | Keep the best-matching locations; resolve ties per molecule |
| Read close to two locations at different edit distances | Observed: for 1.06% of eligible reads with a location within the limits, the emitted location is not a best match; its edit distance is greater in 99.9861% of these and never smaller | The location with fewer edits |
| Main case | In 93.38% of those reads the row barcodes overlap: Space Ranger's BC2 is the 14-base oligonucleotide of the adjacent row, the best-matching location's 15-base BC2 without its first base. In the reads of Supplemental Table S4, the emitted location's 14-base BC2 needs an inserted base (two edits in total) | In the reads of Supplemental Table S4, the location whose 15-base BC2 matches exactly (one edit in total) |
| Read close to two locations at equal edit distance (22.7% of reads with a location within the limits) | One location or none; the emitted location is one of the tied locations for 92.6% of these reads | All best-matching locations kept until molecule resolution |
| A location lies within 2 edits per half but none is emitted | 2.92% of eligible reads with a location within the limits, from 0.001% of reads one edit from the best-matching location to 44.6% of reads four edits from it | No read dropped at decoding |
| A location is emitted but none lies within 2 edits per half | 92 reads | No location assigned |

**Supplemental Table S6. Where the molecules STARgrid counts beyond Space Ranger come from, on two slides.** $M$: the molecules that Space Ranger counted (Space Ranger 4.0.1 on slide H1-VM2JXXK/A1, on the genes of its probe set; 4.1.0 with introns included on the ovarian dataset). $T$: STARgrid's total. $B$ and $H$ count the reads that Space Ranger mapped to a single gene and gave a UMI, with Space Ranger's gene and the raw UMI, at Space Ranger's locations ($B$) or at the locations STARgrid chooses ($H$), under the UMI correction of the matrices. Columns of the lower rows: $D = T - M$; placement $H - B$; gene assignment $T - H$ (STARgrid's genes, selection of reads and handling of UMIs with reads for two genes); deduplication $B - M$. The three components add up to $D$.

| | $M$ / $D$ | $T$ / $H - B$ | $B$ / $T - H$ | $H$ / $B - M$ |
|---|---|---|---|---|
| Colorectal (Flex), H1-VM2JXXK/A1 | 272,612,275 | 279,777,220 | 273,110,393 | 280,257,120 |
| molecules | +7,164,945 | +7,146,727 | −479,900 | +498,118 |
| % of $D$ | 100 | +99.7 | −6.7 | +7.0 |
| Ovarian dataset (RNA-seq), H1-YQJBZ7X/D1 | 187,552,358 | 201,888,275 | 189,504,493 | 198,231,870 |
| molecules | +14,335,917 | +8,727,377 | +3,656,405 | +1,952,135 |
| % of $D$ | 100 | +60.9 | +25.5 | +13.6 |

**Supplemental Table S7. Nucleus by nucleus: paired depth in place and against a far-shift background.** For each nucleus with at least 4 capture locations in the tissue, the paired depth is one minus the mean count on its locations divided by the mean on the same locations moved to each of the 8 neighbouring shifts, one location away (neighbours), or to each of the 16 next shifts out, two locations away (next out). In place and Background are medians in %, and In place − background is their difference in percentage points, each with a percentile 95% interval from 200 draws of spatial blocks of the slide. In place: the nuclei where the registration puts them, median over nuclei. Background: the same statistic with every nucleus first moved 25 capture locations (50 μm) on each axis, along each of the four diagonals, median over the four placements of every nucleus; a moved nucleus can overlap other nuclei, so this is a background placement, not a nucleus-free one. Nuclei: nuclei in place after leaving out those that would cross the edge of the capture area at one of these shifts and those whose mean over these shifts is zero. Deposited map: the nucleus map deposited with the dataset. Our Cellpose: our masks of the ovarian dataset moved by (−3, −3) locations, the dip under our registration. Xenium: the nuclei imaged by Xenium on the same section as the Visium HD run, in four cores of one slide.

| Case | Nuclei from | Nuclei | Against | In place | Background | In place − background |
|---|---|---|---|---|---|---|
| Ovarian | deposited map | 216,104 | neighbours | 4.00 [3.94, 4.09] | 0.63 [0.60, 0.65] | 3.37 [3.31, 3.46] |
| | | | next out | 9.62 [9.45, 9.79] | 1.87 [1.83, 1.92] | 7.75 [7.58, 7.91] |
| Ovarian | our Cellpose | 160,309 | neighbours | 3.04 [2.97, 3.12] | 0.46 [0.44, 0.48] | 2.59 [2.52, 2.67] |
| | | | next out | 8.03 [7.84, 8.21] | 1.46 [1.41, 1.50] | 6.58 [6.39, 6.74] |
| Colorectal | deposited map | 273,394 | neighbours | 4.57 [4.41, 4.74] | 0.87 [0.83, 0.90] | 3.70 [3.55, 3.86] |
| | | | next out | 9.45 [9.19, 9.73] | 2.45 [2.39, 2.53] | 7.00 [6.74, 7.24] |
| Colorectal, H1-VM2JXXK/A1 | deposited map | 192,523 | neighbours | 5.58 [5.44, 5.72] | 1.42 [1.37, 1.49] | 4.15 [4.01, 4.33] |
| | | | next out | 9.88 [9.61, 10.17] | 3.52 [3.38, 3.64] | 6.36 [6.05, 6.74] |
| Lung core VUILD49LA | Xenium | 32,271 | neighbours | 5.38 [4.76, 5.88] | 3.11 [3.03, 3.45] | 2.26 [1.60, 2.85] |
| | | | next out | 9.43 [8.57, 10.36] | 8.05 [7.69, 8.57] | 1.39 [0.46, 2.58] |
| Lung core TILD113LA | Xenium | 18,178 | neighbours | 5.08 [4.59, 5.68] | 3.03 [2.54, 3.19] | 2.05 [1.56, 2.94] |
| | | | next out | 9.43 [8.43, 10.62] | 7.69 [7.25, 8.20] | 1.74 [0.53, 2.99] |
| Lung core TILD111LA | Xenium | 22,708 | neighbours | 3.70 [3.03, 4.26] | 3.03 [3.03, 3.45] | 0.67 [−0.21, 1.20] |
| | | | next out | 6.36 [5.34, 7.34] | 7.69 [7.27, 8.16] | −1.34 [−2.42, −0.33] |
| Lung core VUHD049 | Xenium | 19,113 | neighbours | 2.61 [2.04, 3.03] | 4.00 [3.45, 4.00] | −1.39 [−1.96, −0.58] |
| | | | next out | 4.99 [4.00, 5.88] | 8.99 [8.57, 9.44] | −4.01 [−5.09, −2.69] |

**Supplemental Table S8. Whole-slide and regional offsets under each registration: the dip and the correlation ratio.** Our nucleus masks were placed on the capture grid by each registration and moved over it one capture location at a time. Shifts are (rows, columns) of capture locations of 2 µm, rows running down the slide. Dip: the position of the deepest dip within 12 locations; Depth: one minus the mean count at the dip divided by the median over its 8 neighbouring shifts. Correlation ratio: the capture locations are grouped by their distance from the edge of the nearest nucleus, in bands one location wide (3 inside the nuclei, 6 outside them, and the rest of the tissue); each count is divided by the mean count of the capture locations in tissue in the $5 \times 5$ square around it; the shift is the highest local maximum of the correlation ratio $\eta^2$ of the divided counts on the groups within 12 locations, higher than at its 8 neighbouring shifts (Supplemental Methods). Offset: the distance of the shift from zero. Cellpose: nuclei segmented with Cellpose-SAM, as STARgrid does; StarDist: nuclei segmented with StarDist, the control on the ovarian dataset and the two colorectal slides. Regions: the capture area divided $5 \times 5$; a region is measured when it holds at least 30,000 nucleus locations in tissue. Usable: not on the edge of the search, and in at least 95% of 200 draws of spatial blocks of the slide the shift found lies within one location of the full-data one. At zero: usable regions whose shift is zero. Largest: the largest regional offset. Both usable: of the regions in which the dip and the correlation ratio are both usable, those where they give the same shift (Equal) and shifts one location apart (One off); in none do they differ by more. Deposited: the registration in the output deposited with the dataset; our run: our Space Ranger run on a subsample of 100,000 reads (Supplemental Methods). For each set of nuclei on a slide, the masks and the count image are the same in every row; only the registration changes. Every whole-slide dip and correlation ratio is usable, and on every whole slide the two give the same shift.

| | | Whole slide, dip | | Whole slide, ratio | | Regions usable | | | | Both usable | |
|---|---|---|---|---|---|---|---|---|---|---|---|
| Nuclei | Registration | Shift | Depth (%) | Shift | Offset (µm) | Dip | Ratio | At zero | Largest (µm) | Equal | One off |
| *Ovarian dataset, H1-YQJBZ7X/D1* | | | | | | | | | | | |
| Cellpose | Ours, from the images | (−3, −3) | 2.3 | (−3, −3) | 8.5 | 21 | 21 | 0 | 10.0 | 20 | 1 |
| StarDist | Ours, from the images | (−3, −3) | 3.0 | (−3, −3) | 8.5 | 21 | 21 | 0 | 10.0 | 20 | 1 |
| Cellpose | Space Ranger 4.0.1, deposited | (0, 0) | 3.0 | (0, 0) | 0.0 | 21 | 21 | 20 | 2.0 | 20 | 1 |
| StarDist | Space Ranger 4.0.1, deposited | (0, 0) | 4.0 | (0, 0) | 0.0 | 20 | 20 | 20 | 0.0 | 20 | 0 |
| *Colorectal slide, H1-GMHFWPH/D1* | | | | | | | | | | | |
| Cellpose | Ours, from the images | (0, 0) | 3.1 | (0, 0) | 0.0 | 21 | 21 | 13 | 2.0 | 19 | 2 |
| StarDist | Ours, from the images | (0, 0) | 3.5 | (0, 0) | 0.0 | 21 | 20 | 13 | 2.0 | 18 | 2 |
| Cellpose | Space Ranger 4.1.0, deposited | (0, 0) | 3.3 | (0, 0) | 0.0 | 21 | 21 | 21 | 0.0 | 20 | 1 |
| StarDist | Space Ranger 4.1.0, deposited | (0, 0) | 3.8 | (0, 0) | 0.0 | 21 | 20 | 20 | 0.0 | 20 | 0 |
| *SPATCH slide, H1-WP2FTZD/D1* | | | | | | | | | | | |
| Cellpose | Ours, from the images | (−5, −3) | 2.0 | (−5, −3) | 11.7 | 22 | 22 | 0 | 15.2 | 20 | 2 |
| Cellpose | Ours, with our fitted correction | (0, 0) | 4.1 | (0, 0) | 0.0 | 22 | 22 | 22 | 0.0 | 21 | 1 |
| Cellpose | Space Ranger 3.0.0, our run | (−5, −2) | 2.4 | (−5, −2) | 10.8 | 22 | 22 | 0 | 12.6 | 18 | 4 |
| Cellpose | Space Ranger 4.1.0, our run | (0, 0) | 2.4 | (0, 0) | 0.0 | 22 | 22 | 7 | 4.5 | 18 | 4 |
| *Colorectal slide H1-VM2JXXK/A1* | | | | | | | | | | | |
| Cellpose | Space Ranger 3.0.0, deposited in GEO | (−1, −2) | 2.7 | (−1, −2) | 4.5 | 13 | 16 | 0 | 6.3 | 11 | 2 |
| StarDist | Space Ranger 3.0.0, deposited in GEO | (−1, −2) | 3.1 | (−1, −2) | 4.5 | 18 | 20 | 0 | 7.2 | 16 | 1 |
| Cellpose | Space Ranger 4.1.0, our run | (0, 0) | 3.0 | (0, 0) | 0.0 | 13 | 16 | 12 | 2.0 | 12 | 1 |
| StarDist | Space Ranger 4.1.0, our run | (0, 0) | 3.8 | (0, 0) | 0.0 | 17 | 20 | 14 | 2.0 | 14 | 2 |

**Supplemental Table S9. The window of the local mean.** The correlation ratio (Supplemental Methods) with each count divided by the mean count of the capture locations in tissue in a square window around it, of the side given (in capture locations; half-width in µm), or not divided (none); in bold, the window STARgrid uses. Shifts: usable shifts, by the rule of Supplemental Table S8. Lung: the 152 cores, whole sections and regions of Supplemental Table S11, with nuclei from the H&E image or imaged by Xenium; At truth: shifts at the truth; 2+ off: two or more locations from it. Tumour slides: the 345 whole slides and regions of the placements of Supplemental Table S8; With dip: those where the dip is also usable; 2+ from dip: of these, shifts two or more locations from the dip's. Public slides: the 69 whole slides and regions of Supplemental Table S10; 2+ from SR: shifts two or more locations from Space Ranger 4's registration.

| | | Lung (152) | | | Tumour slides (345) | | | Public slides (69) | |
|---|---|---|---|---|---|---|---|---|---|
| Window | Half-width (µm) | Shifts | At truth | 2+ off | Shifts | With dip | 2+ from dip | Shifts | 2+ from SR |
| none | — | 106 | 75 | 4 | 73 | 62 | 10 | 40 | 6 |
| $3 \times 3$ | 2 | 101 | 81 | 0 | 323 | 310 | 0 | 56 | 3 |
| **$5 \times 5$** | **4** | **137** | **109** | **1** | **341** | **328** | **0** | **64** | **0** |
| $7 \times 7$ | 6 | 136 | 109 | 1 | 343 | 330 | 0 | 66 | 1 |
| $9 \times 9$ | 8 | 136 | 107 | 1 | 341 | 330 | 0 | 68 | 1 |
| $11 \times 11$ | 10 | 135 | 108 | 1 | 339 | 328 | 0 | 69 | 1 |
| $15 \times 15$ | 14 | 131 | 101 | 2 | 320 | 310 | 0 | 69 | 1 |
| $21 \times 21$ | 20 | 131 | 102 | 1 | 300 | 290 | 3 | 68 | 1 |
| $31 \times 31$ | 30 | 131 | 100 | 0 | 253 | 242 | 12 | 67 | 1 |
| $41 \times 41$ | 40 | 131 | 98 | 1 | 199 | 186 | 17 | 64 | 1 |
| $61 \times 61$ | 60 | 130 | 95 | 1 | 171 | 158 | 22 | 61 | 2 |

**Supplemental Table S10. The dip and the correlation ratio on four public Visium HD datasets processed with Space Ranger 4.0.1.** Nuclei: the nucleus map deposited with each dataset, placed by the deposited registration; counts: the deposited 2 µm counts in tissue. Dip: the deepest dip within 12 locations of the whole slide, in (rows, columns) of capture locations of 2 µm; depth is one minus the mean count at the dip divided by the median over its 8 neighbouring shifts, in %. Correlation ratio and Usable: as in Supplemental Table S8. Regions: of the $5 \times 5$ regions holding at least 30,000 nucleus locations in tissue, those where the dip, or the correlation ratio, is usable; At zero and One off: usable regional ratios at zero shift, Space Ranger's registration, and one location from it; none is farther. No dip: of the regions with no dip at all, those where the correlation ratio is usable (—, every region has a dip). Inside, Outside: mean count per capture location in tissue inside nuclei (all locations of nuclei) and 0 to 2 µm outside the nucleus edge. Every whole-slide correlation ratio is usable.

| | Whole slide, dip | | | Ratio | Regions usable | | | | | Mean count | |
|---|---|---|---|---|---|---|---|---|---|---|---|
| Tissue | Shift | Depth (%) | Usable | Shift | Dip | Ratio | At zero | One off | No dip | Inside | Outside |
| human lymph node FFPE | (0, 0) | 7.1 | yes | (0, 0) | 10 of 10 | 9 of 10 | 7 | 2 | — | 9.1 | 10.5 |
| human prostate cancer FFPE | (0, 0) | 2.7 | yes | (−1, 1) | 14 of 23 | 19 of 23 | 6 | 13 | 5 of 6 | 6.8 | 6.2 |
| human kidney FFPE | no dip | — | no | (0, 0) | 2 of 22 | 22 of 22 | 15 | 7 | 1 of 1 | 9.8 | 9.0 |
| mouse brain FFPE | no dip | — | no | (0, −1) | 0 of 10 | 10 of 10 | 2 | 8 | 3 of 3 | 30.7 | 24.1 |

**Supplemental Table S11. Lung tissue measured by Xenium and then Visium HD on the same section.** Tissue: the four cores of one slide (H1-K3WKZHN/D1) and the whole sections of two further slides (identifiers given). Each tissue has two rows for the nuclei segmented from the H&E image: Cellpose-SAM, as STARgrid does, and StarDist, the control; the Xenium nuclei are the same in both rows and their results are given in the first. Registration: Space Ranger 3.0.0 in all of them; by hand for the four cores and, in Loupe Browser 8.0, for H1-JGHRVB7/D1. Mask displacement: the shift that brings the nuclei segmented from the H&E image onto the nuclei imaged by Xenium, measured from the two masks alone, without counts, for the segmentation of the row; this is the truth for the H&E nuclei of the row, and zero is the truth for the Xenium nuclei. Dip: the deepest dip within 12 locations in the counts deposited with the dataset. Correlation ratio and Usable: as in Supplemental Table S8. Within one: the percentage of draws of spatial blocks in which the shift found lies within one location of the full-data one. Shifts not usable are in italics. Shifts are (rows, columns) in capture locations of 2 µm, rows running down the slide. Lower part, regions (the $3 \times 3$ division of each core, the $5 \times 5$ division of a section; regions holding at least 30,000 nucleus locations in tissue): the truth of a region is the mask displacement of that region for the H&E nuclei and zero for the Xenium nuclei. Usable dips and usable ratios: regions where the dip or the correlation ratio is usable, by the distance of its shift from the truth. No usable dip lies two or more locations from its truth.

| | Mask displacement | | H&E nuclei, dip | | H&E nuclei, ratio | | Xenium nuclei, dip | | Xenium nuclei, ratio | |
|---|---|---|---|---|---|---|---|---|---|---|
| Tissue | Shift | µm | Shift | Within one (%) | Shift | Within one (%) | Shift | Within one (%) | Shift | Within one (%) |
| VUILD49LA, Cellpose-SAM | (−2, −3) | 7.2 | (−2, −4) | 100.0 | (−2, −4) | 100.0 | (0, 0) | 100.0 | (0, 0) | 100.0 |
| VUILD49LA, StarDist (control) | (−2, −3) | 7.2 | (−2, −3) | 100.0 | (−2, −3) | 100.0 | Xenium nuclei: as in the row above | | | |
| TILD113LA, Cellpose-SAM | (−4, −2) | 8.9 | (−4, −2) | 100.0 | (−4, −2) | 100.0 | (0, 0) | 100.0 | (0, 0) | 100.0 |
| TILD113LA, StarDist (control) | (−4, −2) | 8.9 | (−4, −2) | 100.0 | (−4, −2) | 100.0 | Xenium nuclei: as in the row above | | | |
| TILD111LA, Cellpose-SAM | (−3, −3) | 8.5 | *(−3, −3)* | 59.0 | (−3, −3) | 100.0 | *(0, 0)* | 88.5 | (0, 0) | 100.0 |
| TILD111LA, StarDist (control) | (−3, −3) | 8.5 | (−3, −3) | 100.0 | (−3, −3) | 100.0 | Xenium nuclei: as in the row above | | | |
| VUHD049, Cellpose-SAM | (−5, −1) | 10.2 | none | — | *(−4, −2)* | 50.5 | none | — | *(−1, −1)* | 68.5 |
| VUHD049, StarDist (control) | (−5, −1) | 10.2 | (−5, −1) | 97.0 | (−5, −1) | 100.0 | Xenium nuclei: as in the row above | | | |
| H1-9QZXXNP/D1, Cellpose-SAM | (−4, −2) | 8.9 | (−5, −2) | 100.0 | (−5, −2) | 100.0 | (0, 0) | 100.0 | (0, 0) | 100.0 |
| H1-9QZXXNP/D1, StarDist (control) | (−4, −2) | 8.9 | *(−4, −2)* | 78.5 | (−4, −2) | 100.0 | Xenium nuclei: as in the row above | | | |
| H1-JGHRVB7/D1, Cellpose-SAM | (−4, −2) | 8.9 | (−4, −2) | 100.0 | (−4, −2) | 100.0 | (0, 0) | 100.0 | (0, 0) | 100.0 |
| H1-JGHRVB7/D1, StarDist (control) | (−4, −2) | 8.9 | *(−4, −2)* | 65.0 | (−4, −2) | 100.0 | Xenium nuclei: as in the row above | | | |

| | H&E nuclei | | | | | | Xenium nuclei | | | |
|---|---|---|---|---|---|---|---|---|---|---|
| | | Usable dips | | Usable ratios | | | | Usable ratios | | |
| Tissue | Regions | At truth | One off | At truth | One off | Two off | Regions | At truth | One off | Two off |
| VUILD49LA, Cellpose-SAM | 0 | 0 | 0 | 0 | 0 | 0 | 5 | 5 | 0 | 0 |
| VUILD49LA, StarDist (control) | 9 | 1 | 2 | 3 | 3 | 0 | | as above | | |
| TILD113LA, Cellpose-SAM | 0 | 0 | 0 | 0 | 0 | 0 | 0 | 0 | 0 | 0 |
| TILD113LA, StarDist (control) | 9 | 5 | 3 | 5 | 3 | 0 | | as above | | |
| TILD111LA, Cellpose-SAM | 0 | 0 | 0 | 0 | 0 | 0 | 0 | 0 | 0 | 0 |
| TILD111LA, StarDist (control) | 8 | 0 | 3 | 3 | 3 | 0 | | as above | | |
| VUHD049, Cellpose-SAM | 0 | 0 | 0 | 0 | 0 | 0 | 0 | 0 | 0 | 0 |
| VUHD049, StarDist (control) | 2 | 0 | 0 | 1 | 0 | 0 | | as above | | |
| H1-9QZXXNP/D1, Cellpose-SAM | 17 | 13 | 0 | 14 | 3 | 0 | 17 | 17 | 0 | 0 |
| H1-9QZXXNP/D1, StarDist (control) | 16 | 2 | 0 | 11 | 3 | 0 | | as above | | |
| H1-JGHRVB7/D1, Cellpose-SAM | 17 | 7 | 3 | 11 | 4 | 0 | 17 | 17 | 0 | 0 |
| H1-JGHRVB7/D1, StarDist (control) | 17 | 2 | 0 | 8 | 6 | 1 | | as above | | |

**Supplemental Table S12. Added counts against shared counts on the H&E image.** The values behind Fig. 6A,B. Shared and added counts, the in-cell fraction and the cell ROC AUC are defined in the Methods; the in-cell fraction expected if counts were placed at random is the share of the area that cells occupy, which differs between slides.

| | Colorectal (Flex) | SPATCH ovarian (Flex) | Ovarian (RNA-seq) |
|---|---|---|---|
| *In-cell fraction* | | | |
| Shared counts | 0.513 | 0.418 | 0.853 |
| Added counts | 0.502 | 0.409 | 0.844 |
| Expected if placed at random | 0.335 | 0.330 | 0.582 |
| *Cell ROC AUC* | | | |
| Shared counts | 0.736 | 0.660 | 0.882 |
| Added counts | 0.678 | 0.587 | 0.857 |

**Supplemental Table S13. Sensitivity of the top-gene comparison with Xenium.** Ovarian dataset against the adjacent-section Xenium run. The first row is the primary comparison reported in the main text: the 20 cancer-panel genes with the most Xenium counts, on 128 μm patches, with Pearson correlation. Each other row changes one of these choices; the variants are reported without adjustment for multiple testing. Genes: the number of genes compared; only 47 panel genes are measured in all count matrices, so the 50-gene variant is the all-gene variant. HARD wins and losses: genes whose correlation with Xenium across patches is higher or lower under HARD than under Space Ranger; no gene was tied and no correlation was undefined. *P*: exact two-sided sign test on the wins and losses. Median correlation: the median over the genes of the correlation of each set of counts with Xenium across patches.

| | | | | | | | Median correlation | |
|---|---|---|---|---|---|---|---|---|
| Variant | Genes | Patch (μm) | Correlation | HARD wins | HARD losses | *P* | Space Ranger | HARD |
| **Primary** | 20 | 128 | Pearson | 15 | 5 | 0.0414 | 0.238 | 0.257 |
| Fewer genes | 10 | 128 | Pearson | 8 | 2 | 0.109 | 0.188 | 0.201 |
| More genes | 30 | 128 | Pearson | 21 | 9 | 0.0428 | 0.218 | 0.210 |
| All eligible genes | 47 | 128 | Pearson | 33 | 14 | 0.00794 | 0.143 | 0.154 |
| Smaller patches | 20 | 64 | Pearson | 13 | 7 | 0.263 | 0.109 | 0.111 |
| Larger patches | 20 | 256 | Pearson | 13 | 7 | 0.263 | 0.430 | 0.454 |
| Rank correlation | 20 | 128 | Spearman | 18 | 2 | 0.000402 | 0.238 | 0.247 |

**Supplemental Table S14. Registration sensitivity on the colorectal slide.** Each entry gives the score with Space Ranger's registration of the microscope image to the CytAssist image, followed by the score with ours, using the same image masks placed through each (Supplemental Fig. S4). The largest change in any STARgrid minus Space Ranger score difference is $1.97 \times 10^{-3}$.

| Counts | In-cell fraction | Cell ROC AUC | Nucleus ROC AUC |
|---|---|---|---|
| Space Ranger | 0.9494 / 0.9483 | 0.8522 / 0.8511 | 0.6980 / 0.7048 |
| STARgrid HARD | 0.9491 / 0.9481 | 0.8549 / 0.8537 | 0.6996 / 0.7064 |
| HARD, STARgrid only | 0.9429 / 0.9417 | 0.6751 / 0.6743 | 0.6133 / 0.6172 |
| HARD, Space Ranger only | 0.9428 / 0.9416 | 0.5983 / 0.5978 | 0.5641 / 0.5655 |
